# Human trust and emotion in the context of collision avoidance with an autonomous mobile robot: An investigation of predictability and smoothness in virtual reality

**DOI:** 10.64898/2025.12.23.696107

**Authors:** Yuta Matsubara, Hideki Tamura, Tetsuto Minami, Shigeki Nakauchi

## Abstract

The integration of mobile robots into human environments requires that they behave in trustworthy ways. Trust develops through interaction, and erratic movements can easily break it. Previous research examined motion predictability and smoothness separately, but their combined effect remains unclear. We investigated how predictability (consistent versus random) and smoothness (gradual versus abrupt) influence human trust and emotion during collision avoidance in virtual reality. Twenty-six participants encountered a robot across repeated trials. We measured subjective valence, arousal, and trust, along with skin conductance responses. Predictability dominated the results. Consistent robot behavior led to increased trust and positive feelings over time. Conversely, unpredictable behavior kept trust low. Smoothness acted mainly as a moderator for arousal; smooth paths reduced the stress caused by unpredictable moves. We also found that prolonged proximity in gradual movements raised physiological arousal, even if users reported feeling calm. These findings suggest predictability drives social acceptance more than smoothness. In order to promote trust, robot designers should prioritize consistent, learnable behaviors.

**Highlights:**

- We examined AMR motion predictability and smoothness in VR collision avoidance.
- Predictable motion significantly improved human trust and valence over time.
- Unpredictable and abrupt behavior increased physiological arousal.
- Smoothness modulated arousal primarily when behavior was unpredictable.
- Predictability is more critical than smoothness for social acceptance of AMRs.

## 1. Introduction

The integration of autonomous mobile robots, i.e., autonomous mobile robots (AMRs), is increasingly transitioning from controlled industrial settings to dynamic environments, including settings that are populated by humans, such as warehouses, hospitals, and restaurants. This integration offers significant benefits in terms of efficiency and service; however, it simultaneously introduces complex challenges to human-robot interactions (HRIs) (Akalin et al., 2022a). If these robots are to function effectively and be accepted, they must not only perform their tasks competently but also behave in ways that humans perceive as trustworthy, comfortable, and socially acceptable. The tasks of establishing and maintaining these elements at an appropriate level represent a key component of this process, as these factors directly influence how humans interact with, rely on, and ultimately benefit from these autonomous systems (Kok and Soh, 2020a).

However, the task of fostering positive human emotions and trust in AMRs involves more than merely ensuring technical reliability (Mara and Meyer, 2022; Firmino de Souza et al., 2025). Psychological factors significantly shape acceptance and perceived trustworthiness, safety, and emotions. Even highly sophisticated autonomous systems can face resistance if their behavior does not align with human expectations or elicits a sense of unease (Li et al., 2024). In the context of HRI, human trust is not a fixed attribute but rather a dynamic construct that evolves over time, including through repeated interactions (Alhaji et al., 2025). For example, the perceived predictability of a robot’s actions and the nature of its movements during navigation can substantially influence a human’s comfort and confidence in the context of interacting with that robot. An AMR that moves erratically or makes sudden, unexpected maneuvers might erode trust and lead to avoidance, regardless of the robot’s underlying safety or efficiency. Conversely, a robot whose actions are understandable and whose movements are perceived as smooth and considerate may elicit a greater sense of security and collaboration.

This study investigates the dynamic interactions between AMR behavior and human subjective experience in a virtual reality (VR) environment. We therefore pose the following questions: How do the predictability and smoothness of an AMR’s avoidance motion jointly influence human trust, valence, arousal, and skin conductance response (SCR) in a collision avoidance scenario that takes place in a VR environment? Specifically, we explore how two key behavioral characteristics of an AMR — the predictability of its evasion path (predictable vs. unpredictable) and the smoothness of its evasive maneuvers (gradual vs. abrupt) — influence participants’ evolving ratings of valence (i.e., positive or negative feeling), arousal (level of activation), and trust rate, alongside their skin conductance response (SCR), which has been identified as an objective measure of arousal, over a series of interactions. In our design, AMR behavior is treated as a potential influence on valence, arousal, and trust. Since SCR is an objective index of arousal, we expect SCR to vary alongside reported arousal under the same behavioral conditions. By examining these subjective responses and objective indices as they evolve over repeated trials performed under different behavioral conditions, we aim to provide insights into how specific robot motion and interaction patterns shape the dynamic nature of human perception and trust in the context of AMRs that are designed to participate in shared human environments. The findings of this research are intended to inform efforts to design more intuitive, trustworthy, and ultimately more effective AMRs for a variety of real-world applications. We investigate AMR behavior as a potential influence on trust and emotional responses. Furthermore, we measure affect both subjectively (EmojiGrid and Likert scale) and physiologically (SCR) and analyze these outcomes.

## 2. Related Works

### 2.1. Emotion

The investigation of subjective emotions during AMR collision avoidance can benefit from the valence-arousal (VA) model. The use of the VA model in the context of HRI can facilitate the comparison of findings across different studies (Spezialetti et al., 2020). Moreover, encounters with an AMR, such as scenarios involving potential collisions, can elicit a wide spectrum of feelings, ranging from slight unease or mild alertness to more pronounced states such as anxiety or even relief. These experiences might not always fit perfectly with predefined discrete emotion labels. The continuous nature of the valence and arousal dimensions thus facilitates a more detailed assessment of these subjective states. Such an assessment can enable researchers to map changes in this effect to specific AMR behaviors, such as variations in the predictability and smoothness of AMRs’ behavior.

Self-report measures are the primary tool used to assess subjective emotional experiences (LeDoux and Hofmann, 2018). In this context, the Self-Assessment Manikin, which was proposed by Bradley and Lang (1994), is a widely used nonverbal pictorial scale. The SAM enables participants to rate their emotional states in terms of valence, arousal, and often dominance, in which context graphical figures are used to represent different levels along a scale. EmojiGrid, which was developed by Toet et al. (2018), is a similar pictorial scale. This measure enables participants to report their emotional state by selecting emojis that are organized on a two-dimensional grid. The horizontal axis of the grid corresponds to valence, whereas the vertical axis corresponds to arousal. This design facilitates the assessment of both dimensions in a single response while preserving the intuitive nature of pictorial representations. Even in VR environments, such efficiency and intuitiveness represent advantages in experimental designs that involve repeated trials, since simplicity and ease of use are important with regard to efforts to minimize participant fatigue and maintain engagement throughout the experiment (Toet et al., 2019).

Physiological measures offer objective reports of emotional processes. Skin conductance response (SCR), which is a key element of electrodermal activity, is a psychophysiological indicator of sympathetic nervous system activity (Boucsein, 2012). When electrodermal activity is measured on the basis of exosomatic methods, which involve passing a small electrical current through the skin, the resulting signal consists of two main components: a tonic level and phasic changes (Braithwaite et al., 2013). On the one hand, the tonic level, which is known as the skin conductance level, reflects slower, sustained variations in skin conductance, thereby indicating baseline arousal or responses to prolonged environmental factors. On the other hand, phasic changes, which are known as skin conductance response, are transient, event-related bursts in skin conductance. SCR reflects changes in skin electrical conductivity that result from variations in sweat gland activity. Since sweat glands are innervated by the sympathetic nervous system, SCR directly measures their activation. These responses, which result from temporary increases in perspiration, can facilitate the observation of individuals’ reactions to specific stimuli. SCR has been recognized as a physiological index that is significantly correlated with arousal levels, irrespective of the eliciting emotion’s positivity or negativity (Ventura-Bort et al., 2022).

With respect to the use of physiological measurements in HRI research, Haney and Liang (2024) conducted a systematic review of emotional and safety responses in industrial environments and analyzed studies that focused on HRI involving an AMRs. Although physiological measurements could facilitate the objective measurement of emotional responses, the analysis conducted by the aforementioned authors revealed that while 70% of the studies that they considered used behavioral assessment methods, only 8% used physiological measurements. Nevertheless, several studies have used SCR to quantify emotional responses in HRI, thus demonstrating that SCR could serve as a reliable measure of emotional arousal in this context (Swangnetr and Kaber, 2013; Bethel et al., 2007; Tiberio et al., 2013).

On the basis of these findings, SCR is used to assess the emotional states linked with general arousal in response to the characteristics and behaviors of robots, such as their movement speed, their proximity to the individual, and the predictability of their actions (Gupta et al., 2024; Apraiz et al., 2025; Rubagotti et al., 2022). The inclusion of SCR measurements in studies of human responses to AMR collision avoidance provides an objective physiological index of arousal. This index can corroborate subjective reports of arousal. Such responses reflect increased physiological alertness or stress, even if subjective reports are influenced by other factors such as social norms or the desire of the individual to appear calm. Nevertheless, importantly, the amplitude of SCR tends to decrease as individuals’ become habituated to repeatedly presented similar stimuli (Juuse et al., 2024). This phenomenon of a decreased response as a result of repeated exposure also applies to individuals’ levels of arousal.

### 2.2. The Human-Centered Concept in Human-Robot Interactions

In the process of considering human-centered principles pertaining to AMRs, researchers occasionally conflate several key concepts or use them interchangeably in terms of the researchers’ behavior. This section aims to clarify the definitions used in this context.

#### 2.2.1. Safety and Perceived Safety

In the context of HRI, the term safety has traditionally been understood to refer to safety from physical harm to humans or property damage as a result of interactions with robots (Akalin et al., 2023). In the case of AMRs, this definition relies on the robot’s capacity to perceive its environment, predict hazards, and perform actions such as those involved collision avoidance without causing harm (S. M. B. P. B. et al., 2025; Wang et al., 2021). In contrast, perceived safety, as suggested by Bartneck et al. (2009), refers to a human’s subjective assessment or feeling of security in the process of interacting with a robot. This concept has been described as “the user’s perception of the level of danger when interacting with a robot and the user’s level of comfort during this interaction”. Although this psychological state is related to objective safety, these two concepts remain distinct. A robot system that meets all engineering safety standards may nevertheless elicit insecurity or anxiety on the part of a user.

Lasota et al. (2017) explored the concept of safety in further detail by distinguishing between physical and psychological safety. According to this extension, the concept of physical safety aims to prevent unintentional human-robot contact that could lead to physical discomfort or injury. In contrast, psychological safety, which is closely connected to perceived safety, focuses on preventing stressful interactions that might cause indirect psychological discomfort or harm. This distinction is important since the harm that can occur in HRI is not solely physical. Interactions that are physically safe but that elicit stress or fear can hinder robot use and negatively affect human well-being. If humans feel unsafe near a robot, irrespective of the actual objective risk entailed by the robot, their productivity could decline; alternatively, humans could alter their work patterns to avoid the robot or refuse to use the technology entirely(Haney and Liang, 2024; Yam et al., 2023).

#### 2.2.2. Trust and Trustworthiness

Mayer et al. (1995) defined trust as “the willingness of a party to be vulnerable to the actions of another party based on the expectation that the other will perform a particular action important to the trustor, irrespective of the ability to monitor or control that other party”. This definition includes three core components: the acceptance of vulnerability, a positive expectation concerning the other party’s future behavior, and an element of risk-taking, which takes the form of ceding some degree of control or relying on the other’s actions. In contrast, Lee and See (2004) defined trust as “the attitude that an agent will help achieve an individual’s goals in a situation characterized by uncertainty and vulnerability”. This definition frames trust as an attitude that mediates reliance, particularly when the corresponding outcomes are uncertain and the individual is vulnerable to the consequences of the agent’s actions. This definition has been widely adopted in studies seeking to examine trust as a behavior (Kraus et al., 2022). While these two definitions are distinct, they are not mutually exclusive. In contrast, they exhibit significant commonalities: “They have in common that there must be some task for whose execution the trustor needs the trustee. If the trustee’s execution is uncertain, the trustor becomes vulnerable. If the trustor accepts this vulnerability, then he/she trusts the trustee (Saßmannshausen et al., 2023).”

These two definitions of trust are also particularly useful for efforts to examine subjective emotions and trustworthiness, even in scenarios such as collision avoidance with a virtual AMR (Kohn et al., 2021). In the context of HRI, trust is generally not a fixed attribute (Souza et al., 2025; Alhaji et al., 2025). Instead, it represents an evolving assessment that is shaped over time through repeated encounters and experiences with the robot. This evolving quality entails that trust becomes stronger or weaker on the basis of an accumulation of evidence pertaining to the robot’s conduct. This characteristic distinguishes trust from concepts such as perceived safety or comfort; namely, trust incorporates a temporal projection that extends from present observations toward future expectations (Kohn et al., 2021; Akalin et al., 2023).

Trust in the context of HRI has evolved from early conceptualizations of automation trust (Malle and Ullman, 2021) to more advanced frameworks that include cognitive, emotional, and behavioral components (Dzindolet et al., 2003). Recent reviews have noted that trust is a multidimensional construct that develops through interactions between human psychological factors and behavioral characteristics of the robot (Campagna and Rehm, 2024; Kohn et al., 2021).

Studies that have examined real-world interactions have led to advancements in our understanding the process by which trust develops. Babel et al. (2022) conducted field studies that demonstrated how environmental context and interaction patterns influence the formation of trust in the context of HRI. These authors reported that trust levels were higher when robots exhibited consistent and predictable behaviors and communication that conveyed information aimed at preventing the robot from being annoying in a public space. These findings suggest that the development of trust in the context of HRI is significantly influenced by the context in which such interactions occur; furthermore, these results suggest that consistency and communication in robot behavior are key factors with respect to the formation of trust. Moreover, these authors also highlighted the bullying of robots by participants in their experiment. Although such behavior focused primarily on ‘testing’ the robot’s capabilities, this finding highlights the potential for negative HRI and reveals distinctions between the trust formation processes that characterize human-human interactions and those that characterize HRI. The findings reported by these authors suggest that the development of trust is susceptible to the robot’s behavioral consistency and its ability to adapt to human preferences and expectations.

Akalin et al. (2022a) used both subjective and objective measures to identify key influences on perceived safety, including trust, comfort, stress, fear, and anxiety (Rubagotti et al., 2022), in the context of HRI. These authors identified comfort, predictability, and trust as major influences on perceived safety, thus improving our understanding of the conditions under which humans feel unsafe.

The development of trust in social robots fundamentally differs from that of trust in human service providers, particularly with respect to the ways in which people attribute responsibility and respond to failures in these contexts. Studies have reported that humans tend to be less forgiving of robot errors than of human errors, and their emotional responses to robot failures are generally more negative (Chen et al., 2021). Additionally, paradoxes such as the uncanny valley effect emerge when robots exhibit highly humanlike emotional capabilities, thereby often leading to decreased trust (Plaks et al., 2022). Recent research has also compared trust dynamics between robotic and human agents, revealing that children may even exhibit a bias that encourages them to trust a reliable robot over an equally reliable human (Stower et al., 2024). Researchers have reported contrasting results regarding how the timing of robotic errors affects trust formation. On the one hand, Rossi et al. (2017) reported that severe errors that occur early in interactions have more detrimental effects on trust than do errors that occur later, thus suggesting that first impressions significantly influence trust formation. On the other hand, Akalin et al. (2022a) reported no significant differences in trust levels between groups that experienced robot failures at the beginning of interactions and groups that experienced such failures at the end of interactions. These authors reported that participants exhibited higher trust ratings in early interactions regardless of error timing. They reported that this finding is potentially to the result of a “positivity bias”, according to which novice users initially tend to trust automation.

In light of the definition and characteristics of trust, the experimental settings that have been designed to study these interactions often focus on conditions of vulnerability for participants, such as those that involve the potential for a virtual collision and uncertainty, particularly when the actions of the AMR are not predictable (Alhaji et al., 2025; Lingam et al., 2025). In such situations, a participant’s main objective is typically to ensure their own safe passage or the completion of a task. Consequently, the trust ratings identified in these contexts directly reflect a participant’s belief in the AMR’s capacity to execute the necessary evasive action or perform its task successfully. The fundamental components of this definition “namely, the attitude indicated by Lee and See (2004), the pursuit of goals, uncertainty, and vulnerability”are thus directly reflected in such experimental arrangements.

In the context of HRI, it is essential to distinguish between trust and trustworthiness (Kok and Soh, 2020b). Trustworthiness is a characteristic of the trustee (robot), whereas trust is a property of the human user (Ashraf et al., 2006). This difference implies that humans may not trust a robot even if it exhibits a sufficiently high level of trustworthiness; conversely, that humans may also trust a robot that is not inherently trustworthy (Kok and Soh, 2020b).

### 2.3. Collision Avoidance Behavior toward AMRs

Given that AMRs that do not account for humans, such as those that cause harm or discomfort to humans via unsafe driving, are difficult for human society to accept, AMRs in human environments are required to operate through space while simultaneously maintaining both physical safety and psychological comfort for the humans whom they encounter. Recent studies have highlighted the importance of focusing on developing navigation and collision avoidance strategies that take social and psychological factors into account (Camara and Fox, 2022).

Several studies have provided insights into human behavioral adaptations that occur in the process of encountering AMRs. Vassallo et al. (2017) examined the natural human avoidance strategies used when the paths taken by participants in the experiment crossed the paths taken by an AMRs, revealing that humans tend to adapt their trajectories at an early point during the interaction stage, which is similar to the case of human-human collision avoidance. Vassallo et al. (2018) also investigated how humans respond to robots that implement strategies that resemble human-inspired interaction rules and reported that participants engaged in more natural collision avoidance behaviors that were similar to human-human interactions when the robots exhibited human-like behavior patterns. Another study revealed that AMRs who exhibited predictable behavior, such as maintaining a straight path or stopping, were perceived as more comfortable than were robots that changed direction. Furthermore, another study indicated that people perceive AMRs that exhibit predictable behaviors, such as maintaining a straight path or coming to a stop, as more comfortable than those that engage in in less predictable movements, such as changing their course (Neggers et al., 2024).

Human actions may also indicate subsequent collision avoidance behavior, even in VR environments. For example, waist rotation has been identified as a key predictor of human avoidance direction(Yamauchi et al., 2025). The experiments conducted by these authors to investigate human-AMR collision avoidance revealed that waist rotation occurs unconsciously before the avoidance movement, thereby serving as an early signal of the intended direction of the human’s movement. Berton et al. (2019) investigated gaze behavior during human-human collision avoidance involving virtual walkers and reported that human gaze patterns are similar between real and virtual environments. In particular, a VR environment encountered through a headmounted display is associated with qualitatively more realistic gaze patterns, thus highlighting the usefulness of collision avoidance experiments conducted in VR environments.

### 2.4. Characteristics of AMR Behavior That Affect Human Perception

#### 2.4.1. Predictability

On the one hand, predictability in the context of HRI refers to the degree to which a robot’s actions conform to a human’s expectations regarding the robot’s behavior in a specific context (Daronnat et al., 2021). On the other hand, legibility pertains to the clarity with which a robot’s motion expresses its underlying intent or goal (Dragan and Srinivasa, 2013). While these two concepts are related, they remain distinct. A motion can be highly legible (i.e., its intent is very clear) but not predictable if that clearly signaled intent conflicts with what the human expected. In the context of this thesis, AMR behaviors included under the ‘predictable’ condition, including consistent collision avoidance behavior, are intended to enhance the robot’s predictability.

In their review, Haney and Liang (2024) reported that the predictability of an AMR’s movement intention has been a common subject of investigation and that design features or cues enhancing predictability have been reported to improve perceived comfort and reduce hesitant behaviors during interactions. Several HRI studies have demonstrated that predictable robot movements generally lead to more positive human experiences, including in terms of increased perceived safety, greater comfort, and higher levels of trust, as well as more favorable emotional responses, such as higher valence and appropriate levels of arousal (Ivanova et al., 2020; Bortot et al., 2013; Akalin et al., 2022b; Haney and Liang, 2024; Rubagotti et al., 2022). Conversely, unpredictable robot movements tend to elicit negative reactions, such as discomfort, surprise, fear, anxiety, and reduced trust. This difficulty in the context of anticipating programmed behavior may explain some of these negative reactions. Kandul et al. (2023) reported that people are significantly less accurate in the context of guessing an AI’s performance than in that of guessing a human’s performance.

#### 2.4.2. Smoothness

Motion smoothness in the context of robotics and HRI generally refers to the continuity and rate of change in a robot’s movement parameters, such as its acceleration, jerk, and curvature (Akinade et al., 2025; Guillén Ruiz et al., 2020). Accordingly, gradual maneuvers, which involve gentle changes in speed and direction, are perceived as smoother than are abrupt maneuvers, which involve rapid or sudden changes. Previous studies have suggested that smoother robot motions are generally preferred by humans (Akinade et al., 2025; Núnez et al., 2016). Such movements are often perceived as more natural, comfortable, and predictable, as well as less threatening, thereby contributing to more positive HRI experiences.

Greenberg et al. (2025), who investigated the effects of a mobile robot’s passing-motion path curvature controlled by Bezier curves in a hallway, reported that such curvature significantly affected the pleasure and arousal of participants. The participants responded more positively to paths that exhibited moderate curvature, thus indicating a degree of smoothness in the turn. In contrast, the participants responded more negatively to paths that were either too sharply curved or excessively straight. These findings suggest a nonlinear, quadratic relationship between path curvature and human emotional responses.

## 3. Research Questions and Hypotheses

### 3.1. Current Gaps and Research Questions

While the individual effects of AMR motion predictability and smoothness on human responses have received attention, a significant gap exists with respect to our understanding of their combined and interactive effects, especially in the context of a safety-critical collision avoidance task. How does the predictability of an AMR’s evasion strategy modulate human responses to its motion smoothness, and vice versa? For instance, is a smooth but unpredictable AMR perceived more negatively than an abrupt but predictable robot? Furthermore, in scenarios that lack explicit visual or auditory cues, researchers can manipulate predictability through the presence or absence of consistent AMR operational patterns. One example of such a pattern is the situation in which the AMR always veers to a participant’s left upon reaching a predetermined distance. Under these circumstances, however, the question of how the interactions between the factors of predictability and smoothness affect key human responses remains largely unanswered. These responses encompass trust and emotions (arousal and valence) as conceptualized within psychological frameworks.

In light of these gaps, we propose the following research question:

- RQ: How do the predictability (predictable vs. unpredictable) and smoothness (gradual vs. abrupt) of an AMR’s motion influence human trust and emotional responses, including in terms of valence, arousal, and SCR, in the context of collision avoidance in a VR environment?

### 3.2. Hypotheses

On the basis of the proposed research question and the literature review, the following hypotheses are proposed:

- H1: A predictable AMR leads to significantly more positive valence ratings, lower levels of arousal (both subjectively reported and as measured via SCR), and higher trust ratings.
- H2: A smoother AMR leads to significantly more positive valence ratings, lower levels of arousal, and higher trust ratings.
- H3: A significant interaction between predictability and smoothness is evident with respect to the valence, arousal, and trust ratings, in which context predictable and smooth AMR motion leads to the most positive responses.

In light of this review of the relevant literature, an AMR that exhibits unpredictable motions is hypothesized to lead to significantly more negative valence ratings, higher levels of arousal (both subjectively reported and as measured via SCR), and lower trust ratings in comparison with the predictable motions condition. Namely, unpredictability in a collision avoidance task directly violates humans’ expectations for controlled, understandable, and safe behavior on the part of an autonomous agent. A smoothly moving AMR is easier to track visually, and its trajectory is less likely to cause a startle response or a collapse of trust, which could otherwise disrupt the human’s comfort and sense of safety.

## 4. Experiment

### 4.1. Overview of the Experiment

We examined how two properties of an autonomous mobile robot (AMR) in the context of collision avoidance, i.e., the predictability of the avoidance motion and the smoothness of the turn, shape human emotion and trust across repeated encounters in VR. The design of this experiment featured two within-subject factors, i.e., “Predictability” (predictable, unpredictable) and “Smoothness” (abrupt, gradual). Valence and arousal were reported on continuous scales. Trust was rated on a seven-point discrete scale. We modeled valence and arousal on the basis of a linear mixed model, i.e., linear mixed model (LMM), and we modeled trust with on the basis of a cumulative LMM, i.e., cumulative link mixed model (CLMM), with the aim of accounting for its ordinal scale. Furthermore, we analyzed the peak of skin conductance response (SCR) amplitude as an objective index of arousal by using a generalized LMM, i.e., generalized linear mixed model (GLMM).

Across all the outcomes, predictability was identified as a key factor. For valence, a significant interaction between predictability and trial number indicated that the predictable motion led to a positive within-block trend, whereas the unpredictable motion did not lead to any change. For arousal, a three-way interaction was significant. Unpredictable-abrupt avoidance motions were associated with an increasing trend, whereas predictable-abrupt and predictable-abrupt avoidance motions were associated with decreasing trends. For trust, the model indicated a main effect of predictability and an interaction effect between predictability and trials. Analysis confirmed that trust increased across trials only under the predictable motion condition. With respect to SCR, the model indicated lower responses under the predicable motion condition and higher responses under the condition involving the gradual turn, and no significant interactions among the factors were observed.

All interactions and slopes are presented in the figures and tables included in the results section. A summary of the raw data and diagnostic checks for the analysis are provided in the Supplement.

### 4.2. Materials and Methods

#### 4.2.1. Participants

The sample size was calculated with the assistance of PANGEA (Westfall, 2015) on the basis of a two-by-two factorial analysis of variance (ANOVA) including the effect size *d* = 0.35 and *α* = 0.05 to ensure the statistical power of 0.8 for the interaction, resulting in a minimum sample size of twentysix. A total of twenty-six healthy adult members of Toyohashi University of Technology (including three females and twenty-three males; mean age = 22.69; standard deviation *SD* = 2.17) participated in this experiment; all of these individuals were right-handed according to the FLANDERS handedness questionnaire (Nicholls et al., 2013; Okubo et al., 2014) (mean score = 9.69; standard deviation *SD* = 1.05) and had normal or corrected-to-normal vision. The ethics committee of Toyohashi University of Technology approved this study, and all the participants provided written informed consent before their participation in this research.

#### 4.2.2. Apparatus and Stimuli

The stimuli were presented in a VR environment through the use of SteamVR (1.24.6) and Unity (2021.3.4f1) software. The participants were equipped with a head-mounted display (HTC VIVE Focus Vision, HTC, Taoyuan, Taiwan) with a resolution of 2448 × 2448 pixels per eye, totaling 4896 × 2448 pixels, and a refresh rate of 90 Hz. The controller was gripped in the participant’s dominant hand and used to respond to the questionnaires that were presented at the end of each trial. See Appendix A for the equipment used in this research. Despite the differences between VR and the real world, experiments in VR environments are valuable for efforts to investigate HRI and human emotions (Greenberg et al., 2025; Li et al., 2019; Shariati et al., 2018; Pauw et al., 2022). Furthermore, VR experiments offer several unique advantages, including more robust and reproducible control over a robot’s motion and the ability to prevent any harm from resulting from physical contact with a physical AMR.

The virtual environment consisted of a 7.0 m × 6.0 m area that included four walls, a floor and a ceiling. The VR environment was designed to resemble a real-world room. The walls were textured with white paint, and the floor was covered with a gray and black checkered pattern. The ceiling was a plain white surface. The starting point of the participant, which was represented by a white square on the floor, was located 2.0 m away from the center. The destination point, which was represented by a white circle on the floor, was located 3.5 m away from the starting point through the center of the area (3.8 m and 1.8 meter, respectively, in the practice trials). Figure 1 presents a schematic diagram of the VR environment.

**Figure 1:**
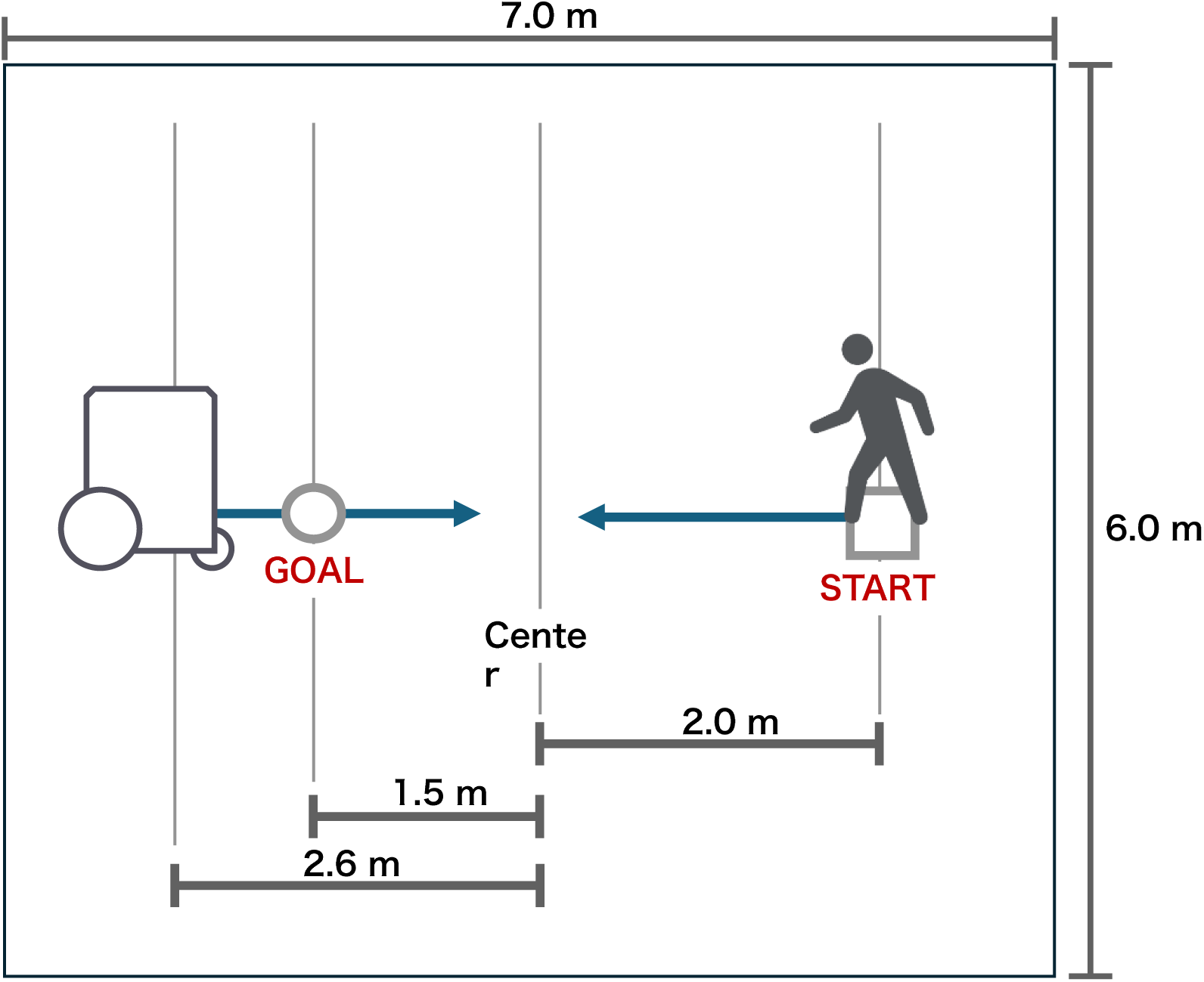
Schematic diagram of the VR environment

The AMR model replicated the MegaRover Ver.2.1 (Vstone, Osaka, Japan) with a custom-made frame of 353 mm × 396 mm × 800 mm (width × depth × height). The AMR moved at a constant speed of 0.5 m*/*s. The AMR was positioned in front of the participant at a distance of 4.6 m; thus, the distances from the center to the starting point were 2.6 m (3.8 m and 1.8 meter, respectively, in the practice trials). The behaviors of the AMR were programmed with C# in Unity. No models were used to represent the participant’s body in the virtual environment.

A cubic Bezier curve was used to control the AMR’s trajectory. Several studies on robot path planning and HRI have employed Bezier curves as a part of their trajectory generation algorithm(Greenberg et al., 2025; Duraklı and Nabiyev, 2022). Bezier curves are parametric curves that are inherently smooth (i.e., they possess continuous derivatives up to a certain order, depending on the degree of the curve) and that offer control over their shape through the manipulation of control points. This approach facilitates the programmatic manipulation of the inflection point in the AMR’s movement. It also enables key parameters such as turning direction and curvature, which lead to differences between the conditions, to be controlled on the basis of the same formula.

#### 4.2.3. Tracking and Measurement

Gaze data during the trial were recorded with the assistance of a VIVE Focus Vision head-mounted display. The gaze data were recorded at a sampling rate of 90 Hz and were used to calculate gaze changes and the corresponding variability.

Three-dimensional data concerning the participants were captured through the use of five VIVE Ultimate Trackers (HTC, Taoyuan, Taiwan) at a sampling rate of 90 Hz. These trackers were attached to the participant’s waist, right wrist, left wrist, right ankle, and left ankle. A VIVE Cosmos Controller (HTC, Taoyuan, Taiwan) was held in the participant’s dominant hand and used both to respond to the questionnaires presented at the end of each trial and to detect collisions with the AMR. Namely, when such a collision occurred, the controller provided vibrotactile feedback.

Participants’ electrodermal activity was recorded with the assistance of a BioRadio (Great Lakes NeuroTechnologies, Cleveland, OH, USA) at a sampling rate of 1000 Hz. Electrodermal activity was measured through the use of three electrodes that were placed on the participant’s index finger, middle finger, and back of the nondominant hand. The BioRadio was connected to a desktop PC via Bluetooth, and the electrodermal activity data were recorded with the assistance of MATLAB R2024b via the BioRadio API. To extract SCR from the electrodermal activity, a 100 Hz low-pass filter for noise removal was used, after which the signal was downsampled to 100 Hz. The data were subsequently smoothed on the basis of a simple moving average with a 50 ms window. For baseline correction, the onset of an auditory cue was designated as time 0, and the average signal from the 0 to 1000 ms interval served as the baseline. The SCR was then extracted from this baseline-corrected electrodermal activity signal. The purposes of collecting SCR data were to obtain a physiological measure of arousal during approach and avoidance, independent of self-reports, and to evaluate the associations between this factor and subjective ratings of arousal. Consequently, participants were equipped with a head-mounted display, a controller, trackers, and a BioRadio.

#### 4.2.4. Experimental Design

The experiment involved two within-subject factors: the predictability of the AMR’s behavior (predictability: predictable or unpredictable) and how the AMR sought to distance itself from the participant (smoothness: abrupt or gradual). Each combination of factors was repeated twelve times, resulting in 48 trials. The experiment consisted of four blocks, including 12 trials per block. Each block was assigned each combination of conditions. The order of the blocks was randomized across participants.

We treated predictability as a cognitive property that referred to the extent to which the timing and the side of the AMR’s avoidance motion were in line with a participant’s expectations regarding what the AMR would do next. Furthermore, we treated smoothness as a kinematic property that focused on the continuity of the avoidance trajectory in terms of curvature. This separation is in line with common practices in the field of HRI, where predictability and motion quality are analyzed on distinct axes. The two factors target different aspects of human perception and are theoretically separable: predictability addresses the performance of the behavior in question, whereas smoothness addresses the manner of movement. For example, a review of industrial HRC considers the antecedents of trust in terms of two axes: performance-related factors (e.g., reliability and predictability) and robot attributes (e.g., proximity and personality) (Simões et al., 2022; van den Brule et al., 2014). van den Brule et al. (2014) manipulated task performance and motion fluency (smooth vs. trembling) and reported that both factors had main effects on trustworthiness, thus facilitating an analysis of performance-related predictability in addition to motion style.

The predictability factor was manipulated by altering the trigger conditions, including the distance at which the avoidance motion was initiated and its direction. Two levels were defined as follows:

- “Predictable” motion: The AMR initiated its avoidance motion at a fixed distance of 2.0 m from the approaching participant. The direction of avoidance was always to the participant’s left.
- “Unpredictable” motion: The trigger distance was randomized for each trial to occur between 0.5 m and 3.0 m. The direction of avoidance was also randomized for each trial, with a 50% probability of moving to the participant’s left and a 50% probability of moving to their right.

The smoothness factor was manipulated by altering the position of the control points for the Bezier curve. Two levels were defined:

- “Gradual” motion: The AMR executed a smoother, more sweeping turn. The control points were positioned to create a wide, gentle arc. This approach resulted in a trajectory with low curvature, in which context the AMR maintained a smaller lateral separation distance from the participant throughout its avoidance motion.
- “Abrupt” motion: The AMR executed a more sudden, sharp turn. The control points were positioned to create a sharp, sudden turn. This approach resulted in a trajectory with high curvature, thus leading to a greater final separation distance from the participant but a more rapid change in the AMR’s direction.

Hence, we manipulated the two factors with disjoint control parameters to generate the AMR’s motion. Predictability was associated with variation in the trigger policy for the avoidance motion. Smoothness was associated with variation in the Bezier control points used for the evasive segment with the aim of creating a low-curvature path or a high-curvature path. By design, preonset trajectories were the same across different smoothness levels and provided no cues regarding when or to which side the AMR would execute the evasive motion. After onset, we applied the same curvature template to either side. Thus, timing and side uncertainty (predictability) and trajectory continuity (smoothness) were controlled by nonoverlapping parameters, thus ensuring that the factors remained orthogonal at the stimulus level.

Figure 2 illustrates the avoidance motion that occurs on the basis of the combination of each condition.

**Figure 2:**
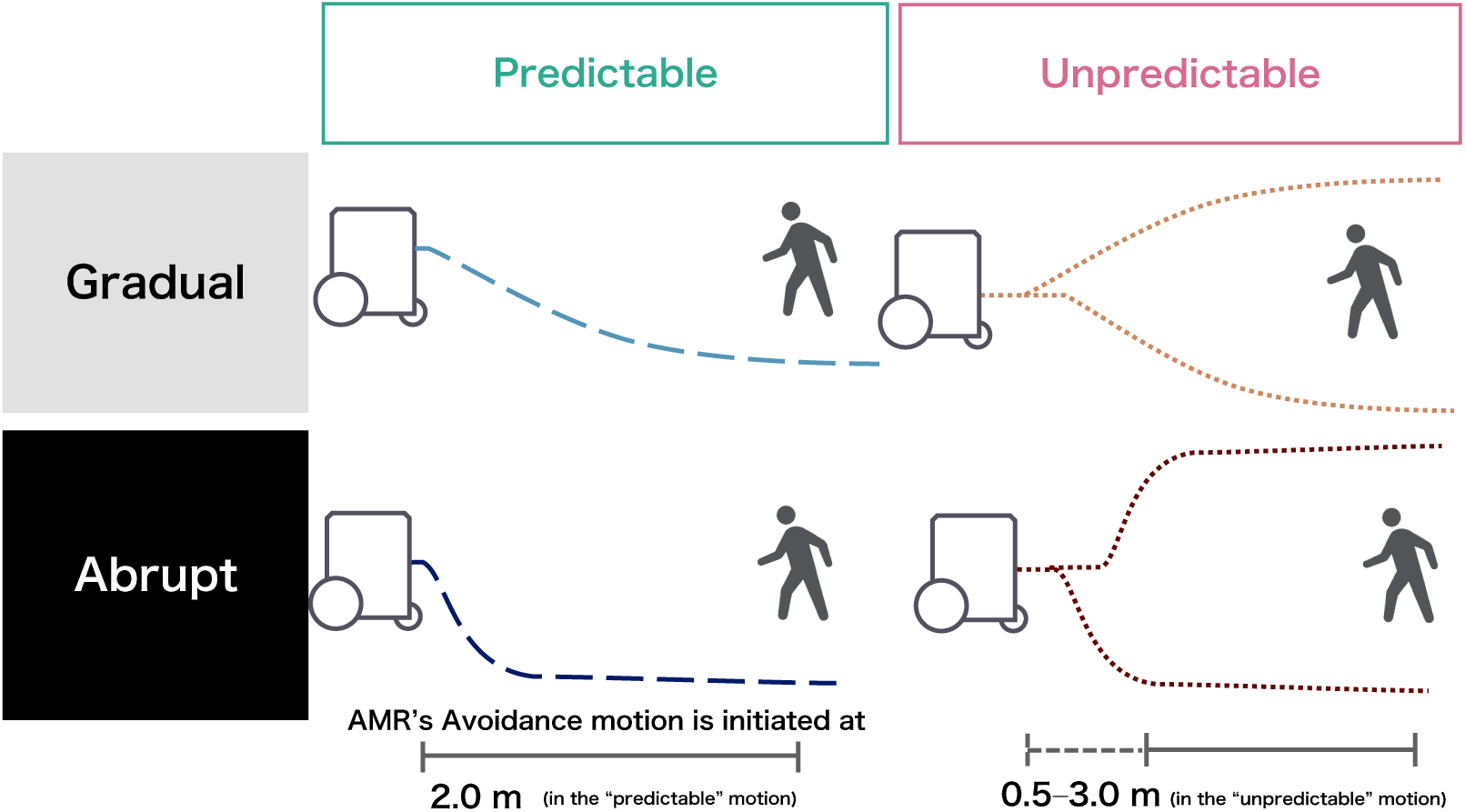
AMR trajectory for each combination of different levels of the conditions experienced by participants.

Many studies have used a single custom question to assess trust in the context of HRI (Souza et al., 2025). Furthermore, the process of evaluating trust through multiple questions after each trial can interrupt the experimental flow and increase the complexity of such research. Therefore, we employed a seven-point Likert scale survey after each trial to measure trust, thereby minimizing participant burden while preserving the accuracy of the measurement.

#### 4.2.5. Procedure

The experiment involved 48 trials, including 12 trials for each of the four experimental conditions. The trials were sorted into four blocks by condition. The participants could take breaks at the end of any trial during the experiment. Furthermore, after two blocks had been completed, the experimenter explicitly asked the participant whether they wished to take a break.

The participant was instructed to move toward the target point at a comfortable pace and to avoid colliding with the AMR if necessary. The AMR executed its avoidance motion as defined by the smoothness and predictability conditions. Figure 3 illustrates the AMR’s behavior as observed by the participant during a trial in the virtual environment.

**Figure 3:**
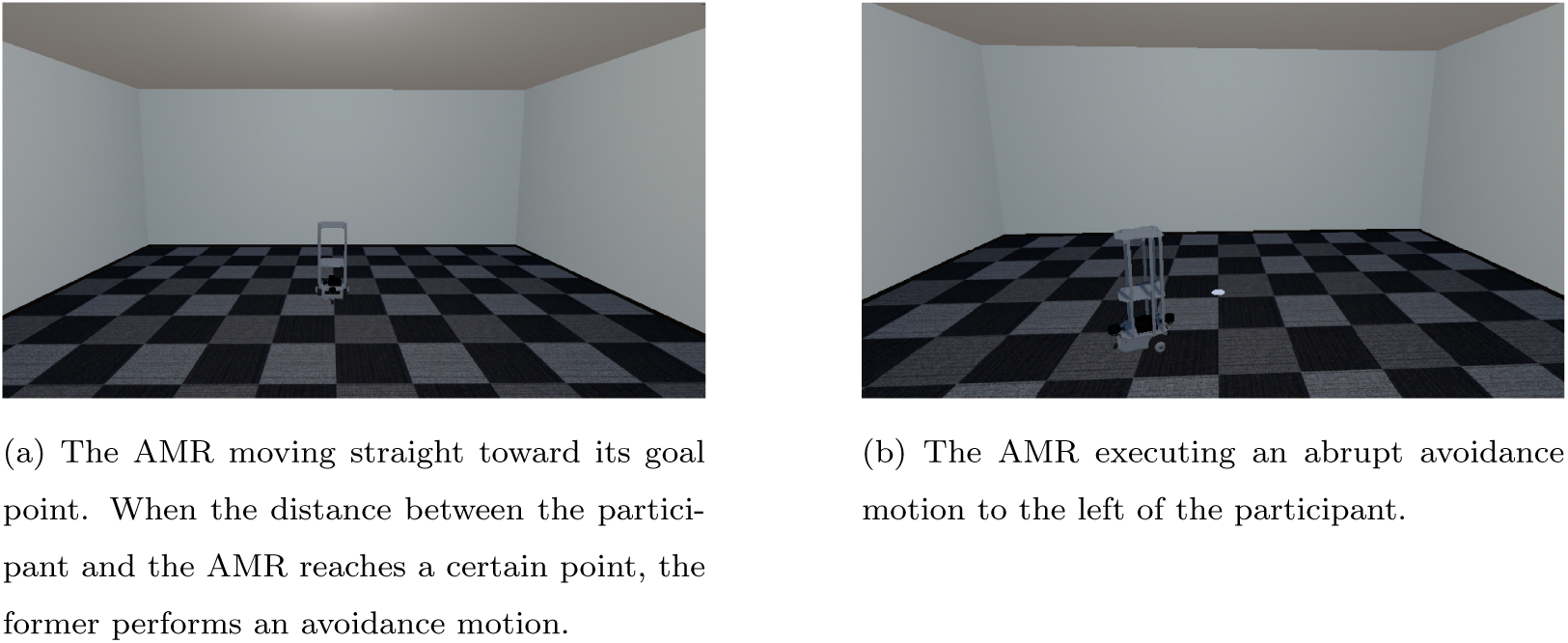
The AMR behavior observed by the participant during a trial in the virtual environment.

The flow of a single trial is illustrated in Figure 4. The trial began with the participant standing at the starting position of the virtual room. The AMR was positioned at a distance of 4.6 m in front of the participant and facing them. Through an auditory cue from the head-mounted display, the participant was instructed to move toward the target point, which was located 3.5 m away from the starting point, at a comfortable and natural pace. When the participant reached the trigger distance (which could be fixed or variable, depending on the predictability condition), the AMR autonomously executed its avoidance motion as defined by the smoothness condition (curvature) and the predictability condition (direction) for the trial in question. Once the participant had reached the target point, the EmojiGrid appeared in front of the participant. The horizontal axis indicated valence, and the vertical axis indicated arousal. Both dimensions were recorded and analyzed as two separate continuous measures. The arrangement of the emoji on the periphery of the EmojiGrid was altered between blocks. These alterations included vertical, horizontal, and combined vertical–horizontal inversions. Thus, emotions represented by certain coordinates changed across blocks. The participants first used a handheld controller to rate their emotional valence and arousal simultaneously. The initial point of the EmojiGrid was randomly selected within the grid for each trial. Subsequently, the participants rated their trust in the AMR that they had just encountered on a 7-point Likert scale, in which context 1 indicated ‘Do not trust at all’ and 7 indicated ‘Trust completely’. The initial point for the Likert scale was randomly selected for each trial. After the trust rating was submitted, the participant was asked to return to the starting point.

**Figure 4:**
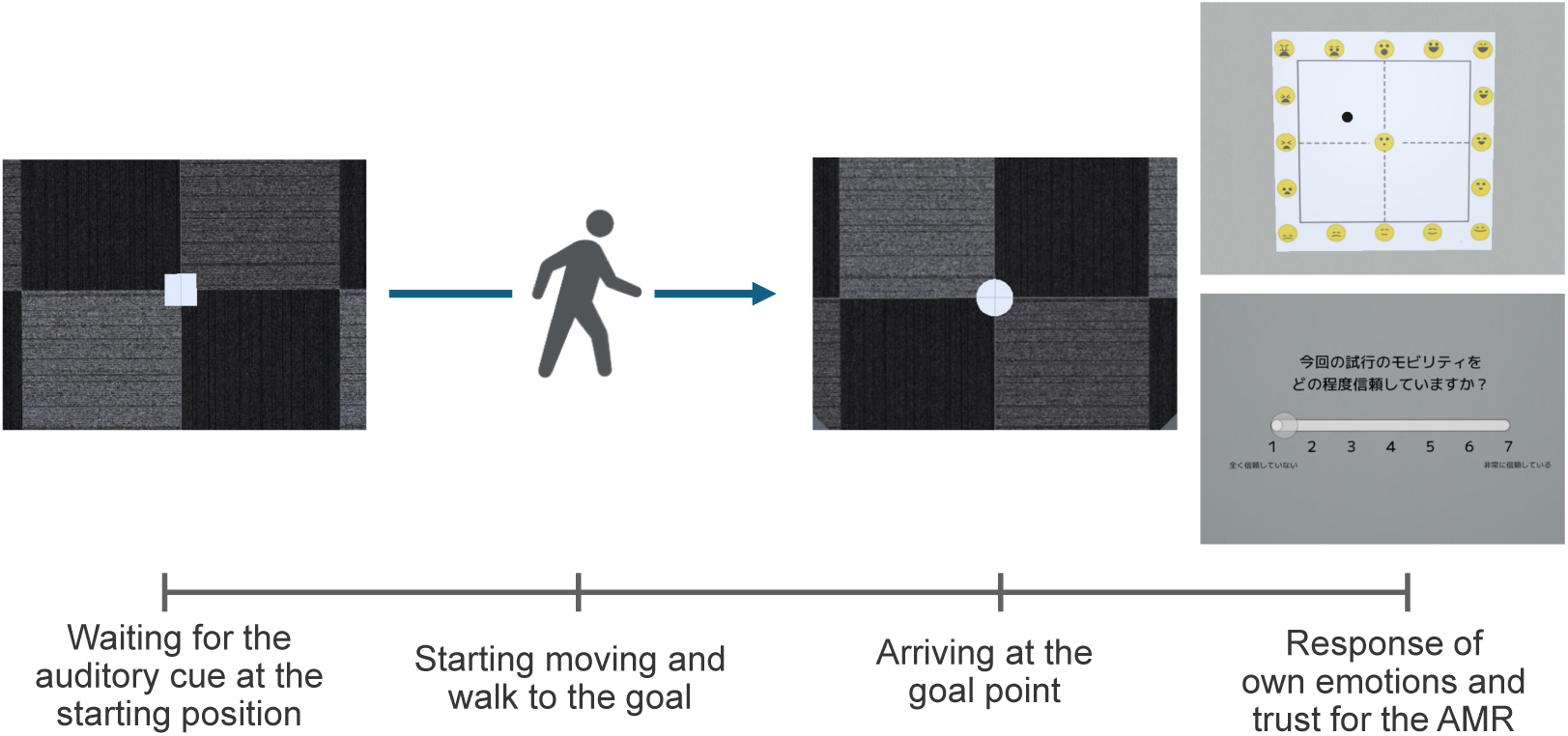
The flow of a single trial in the experiment.

Before the main trials began, the participants experienced three practice trials to familiarize themselves with the experimental procedure. In the practice trials, the AMR did not execute any avoidance motion. Following the completion of all 48 trials, the participant removed the head-mounted display and sensors. The experimenter subsequently collected qualitative feedback, including participants’ impressions of the experiment, through a verbal debriefing session.

#### 4.2.6. Data Analysis

The data thus obtained were processed and analyzed with the assistance of R 4.4.1 software. In the experiment, the analysis was conducted for the period between the presentation of the auditory signal, which represented the start of the trial, and the moment when the participant reached the goal position. This time frame was normalized on a percentage scale.

For each subjective measure, we aimed to understand the influence of the experimental manipulations, i.e., predictability (predictable vs. unpredictable) and smoothness of the evasion (gradual vs. abrupt), as well as the progression of trials within each experimental block. Although the initial power analysis and design were based on a 2×2 factorial ANOVA, all subjective ratings were analyzed on the basis of a linear mixed models (LMMs) with the goal of determining how the temporal evolution of subjective ratings differed across the four experimental conditions (i.e., predictable-gradual, predictable-abrupt, unpredictable-gradual, unpredictable-abrupt). For these models, the participant ID served as a random effect to account for individual differences in the baseline responses. Fixed effects included predictability, smoothness, the centered within-block trial number (Trial in Block centered), and all the two-way and three-way interactions among these factors.

With respect to valence and arousal, which were treated as continuous variables through the use of EmojiGrid, an LMMs involving the lmer() function (provided by lme4 v1.1.35.5; (Bates et al., 2015)) was employed; this approach entails certain assumptions, including that of a Gaussian error distribution. With regard to the trust rate, which was scored on a 7-point Likert scale (ranging from 1 to 7), a cumulative link mixed model (CLMM) involving the clmm() function (provided by ordinal v2023.12.4.1; (Christensen, 2023)) was considered to be more appropriate. This approach was chosen because CLMMs explicitly models the ordinal nature of the data, thereby preserving the ranked order of the categories without assuming equal intervals between each set of categories, which represents a limitation in situations in which ordinal data are treated as continuous in a LMM. The CLMM used a probit link function.

Model selection for each dependent variable involved comparing the candidate models with a variety of fixed and random effect structures using relevant indices, including Akaike’s information criterion, the Bayesian Information Criterion, r-squared, adjusted r-squared, and root-mean-square error, in line with the compare performance() function (provided by performance v0.13.0; (Lüdecke et al., 2021))(Akaike, 1987; Neath and Cavanaugh, 2012). The random effect structure was determined first, followed by the fixed effects. We performed diagnostic checks on all final models by using the check model() function (provided by performance v0.13.0) to assess assumptions such as those pertaining to the normality and homogeneity of residuals, as well as the normality of random effects. Significant interactions were explored in further detail on the basis of simple effects analyses and interaction plots that were generated with the assistance of the emmeans() function and emmip() function (provided by emmeans v0.13.0; (Lenth, 2024)). Significance levels were set at *α* = .05.

We analyzed SCR with the goals of testing our arousal hypotheses (regarding the effects of predictability and smoothness) and assessing convergence with subjective arousal. The primary outcome was peak SCR amplitude within the normalized trial window. As recommended by Lonsdorf et al. (2019), we established a minimum response criterion of 0.05 µS. Participants for whom 66% or more of their trials fell below this threshold were excluded from the analysis, which resulted in the removal of two individuals from the SCR amplitude analysis. Additionally, we excluded one trial in which a participant reported that an electrode had become detached during the experiment.

## 5. Results

### 5.1. Subjective Survey

#### 5.1.1. Valence

A LMM was fitted to the valence data. The final model structure was determined as follows: the model that exhibited the best fit through the model selection process described in the Data Analysis section was Valence ∼ Predictability * Smoothness * Trial_in_Block_centered + (1 *|* ID). See Appendix B for distributions of the ratings (Figure S2 and S3) and Appendix C for diagnostic checks (Figure S14).

Table 1 presents a summary of fixed effects. The LMM revealed a significant two-way interaction between predictability and the centered within-block trials (Estimate = 0.022, SE = 0.01, *t*(1222) = 2.075, *p* = 0.038). This interaction indicates that the valence changed across trials in a manner that depended on whether the robot’s motion was predictable or unpredictable. No other two-way interactions (predictability × smoothness: *p* = 0.147, smoothness × centered within-block trials: *p* = 0.928) or threeway interactions (predictability × smoothness × centered within-block trials: *p* = 0.261) reached the level of significance.

**Table 1:**
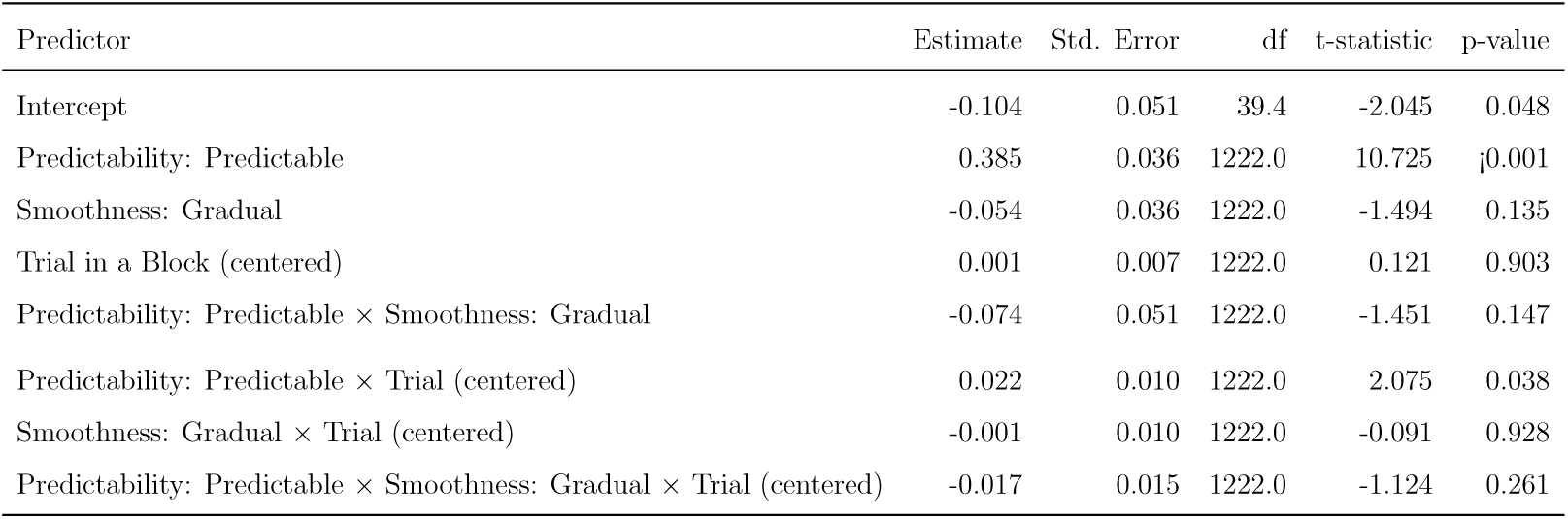
Fixed-effects coefficients of the LMM for valence. The model includes predictability, smoothness, and centered within-block trials as fixed effects, as well as ID as a random effect.

We conducted a simple slope analysis to decompose the significant interaction between probability and the centered within-block trials. This analysis separately estimated the slope of centered within-block trials, i.e., the approximate rate of change in valence per trial, for the predictable and unpredictable conditions; this rate was averaged across smoothness conditions as a result of the nonsignificant higher-order interactions involving smoothness and trial. The results are presented in Table 2. On one hand, valence significantly increased over trials within a block when evasions were predictable (slope = 0.014, SE = 0.005, *t*(1229.04) = 2.634, *p* = 0.009). On the other hand, when the AMR’s collision avoidance behavior was unpredictable, the change in valence across the trials was not significantly different from zero (slope = 0, SE = 0.005, *t*(1229.04) = 0.081, *p* = 0.936). The difference between these two slopes was not statistically significant (*p* = 0.071, Tukey adjusted). Figure 5 illustrates this significant interaction, thereby highlighting a positive trend for valence in the predictable condition.

**Figure 5:**
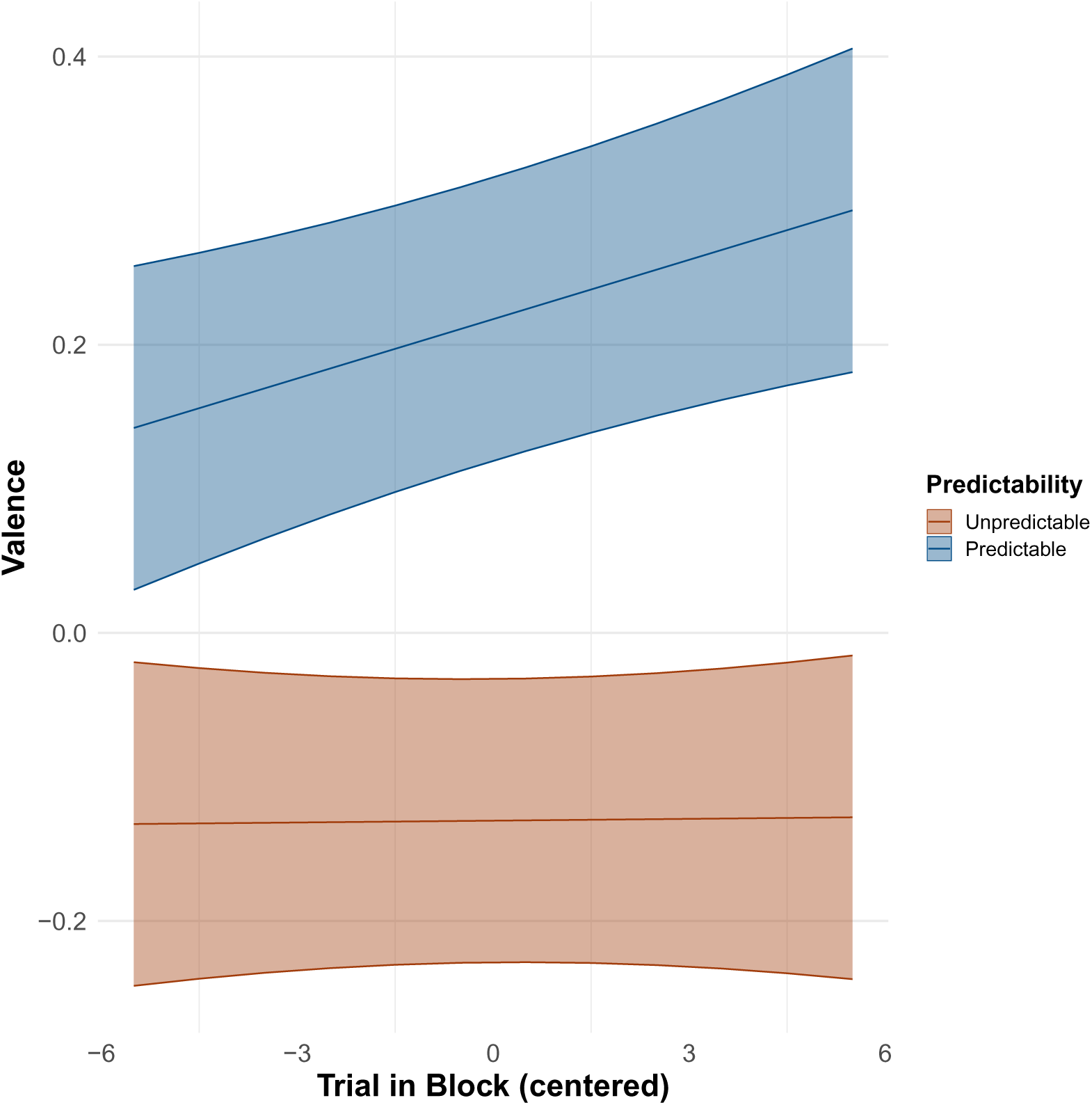
Interaction plot of the best model for valence. The x-axis indicates the centered within-block trials, in which context -5.5 represents the first trial in a block and 5.5 represents the last trial. The y-axis indicates the estimated valence rating on its original scale. The red line and area represent unpredictable conditions, whereas the blue line and area represent predictable conditions. The shaded area represents the 95% confidence intervals.

**Table 2:** Simple slopes of centered within-block trials for each of the four conditions.

| Level of Predictability | Trend of Trial in a Block | SE | 95% CI | t-statistic | p-value |
| --- | --- | --- | --- | --- | --- |
| Unpredictable | 0.000 | 0.005 | [-0.010,0.011] | 0.081 | 0.936 |
| Predictable | 0.014 | 0.005 | [0.004,0.024] | 2.634 | 0.009 |

#### 5.1.2. Arousal

A LMM was also fitted to the arousal data. The final model structure was as follows: Arousal ∼ Predictability * Smoothness * Trial_in_Block_centered + (1 + Trial_in_Block_centered *||* ID). See Appendix B for distributions of the ratings (Figure S4 and S5) and Appendix C for diagnostic checks (Figure S15).

Table 3 presents a summary of fixed effects. The LMM for arousal indicated a significant three-way interaction among predictability, smoothness, and the centered within-block trials (Estimate = 0.046, SE = 0.015, *t*(1196) = 3.143, *p* = 0.002). This three-way interaction indicated that the change in arousal over different trials within a block depended on whether the specific combination of predictability and smoothness that characterized the evasive movement was predictable or unpredictable.

**Table 3:**
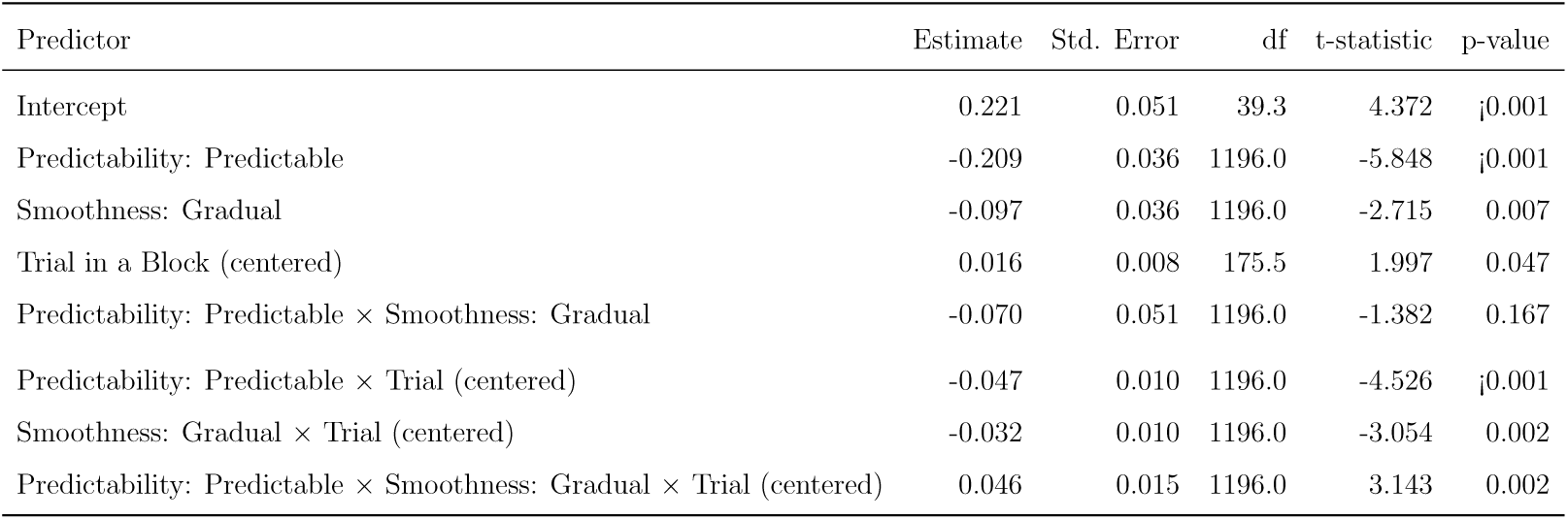
Fixed-effects coefficients of the LMM for arousal. The model includes predictability, smoothness, and the centered within-block trials as fixed effects, alongside ID and the centered within-block trials as a random effect.

To analyze the significant three-way interaction, we conducted a simple slope analysis to examine the slope of the centered within-block trials for each of the four conditions. The results are presented in Table 4. In the unpredictable-abrupt condition, arousal increased significantly as the trials progressed (slope = 0.016, SE = 0.008, *t*(182.149) = 1.979, *p* = 0.049). In the predictable-abrupt condition, arousal decreased significantly across the trials (slope = -0.031, SE = 0.008, *t*(182.149) = -3.818, *p <*0.001). In the unpredictable-gradual condition, the change in arousal over the trials did not reach the level of statistical significance (slope = -0.016, SE = 0.008, *t*(182.149) = -1.933, *p* = 0.055). In the predictable-gradual condition, arousal decreased significantly across the trials (slope = -0.016, SE = 0.008, *t*(182.149) = -2.037, *p* = 0.043).

**Table 4:**
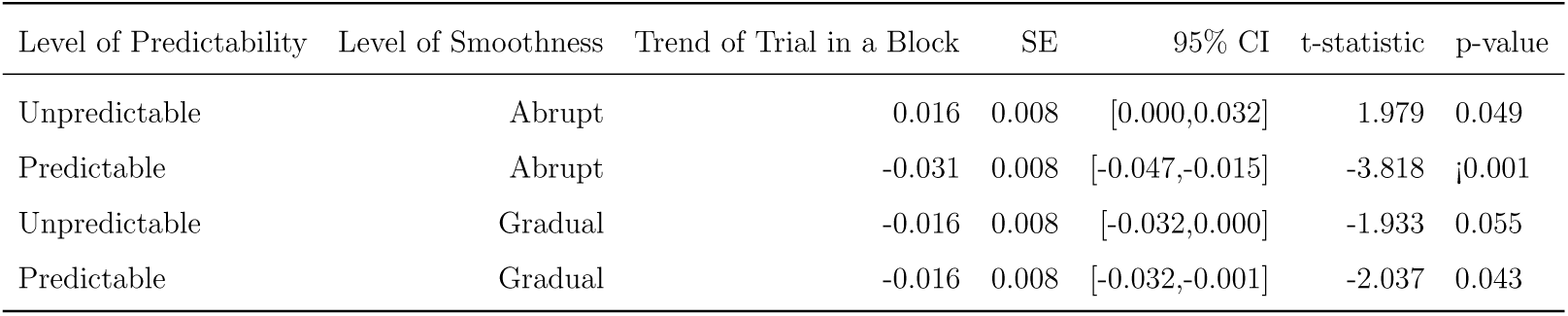
Simple slopes of the centered within-block trials for each of the four conditions.

Pairwise comparisons of these slopes indicated significant differences between the unpredictable-abrupt condition and the other conditions (vs. predictable-abrupt condition:*p* = 0, vs. unpredictable-gradual condition:*p* = 0.013, vs. predictable-gradual condition:*p* = 0.01, Tukey adjusted). Figure 6 represents these trends in arousal across the conditions over the different trials.

**Figure 6:**
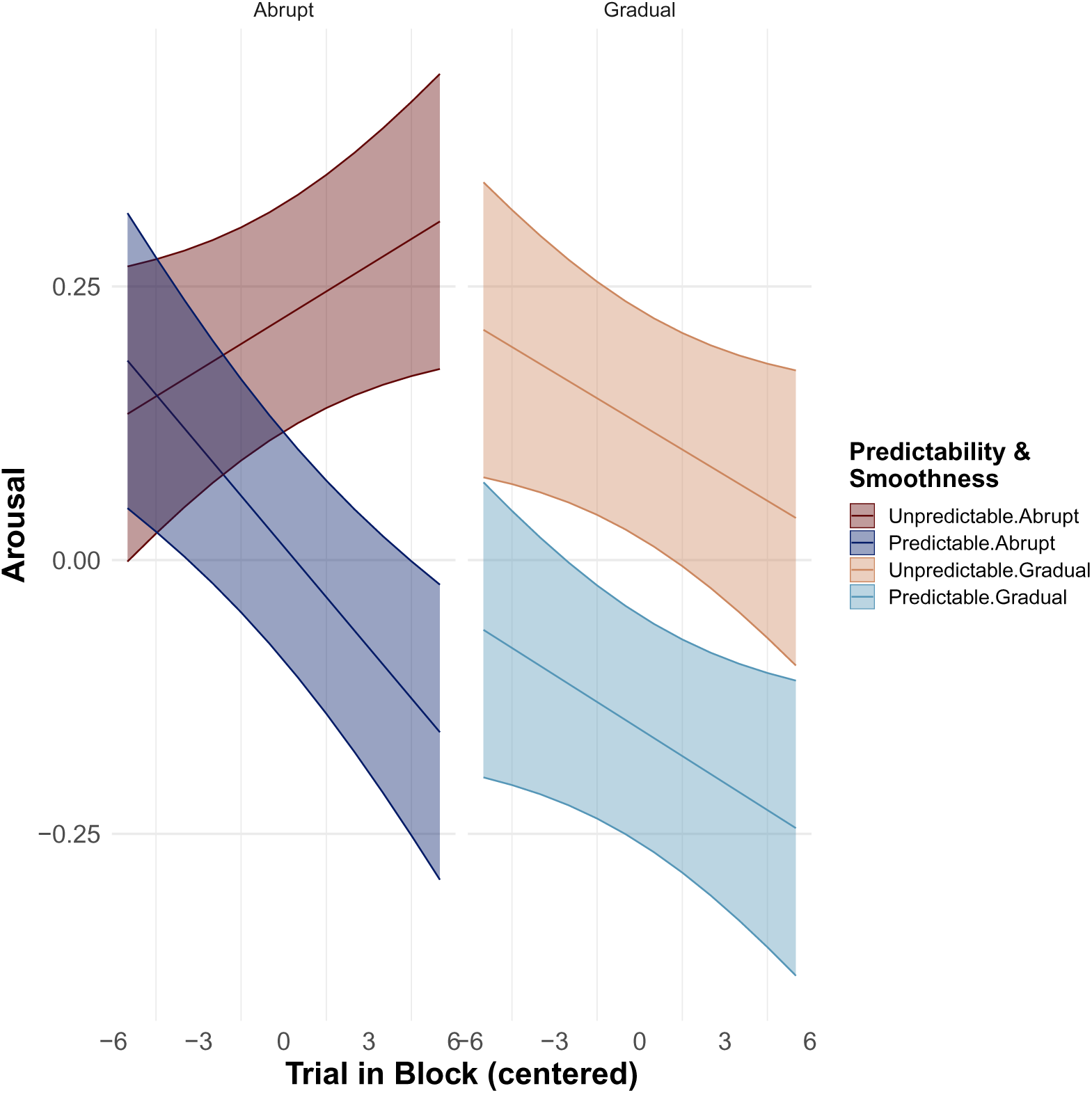
Interaction plot of the best model for arousal. The x-axis indicates the centered within-block trials, in which context -5.5 represents the first trial in a block and 5.5 represents the last trial. The y-axis indicates the estimated arousal rating on its original scale. The dark red line and area represent the unpredictable*abrupt conditions, the dark blue line and area represent the predictable*abrupt conditions, the light red line and area represent the unpredictable*gradual conditions, and the light red line and area represent the unpredictable*gradual conditions. The shaded area represents the 95% confidence intervals.

#### 5.1.3. Trust Rate

We examined participants’ self-reported trust rate, an ordinal measure on a scale ranging from 1 to 7, to identify their degree of trust in the robot. Given the ordinal nature of the trust rate data, we employed a cumulative link mixed model (CLMM) with a probit link function. The final model structure was trustRate ∼ Predictability * Smoothness * Trial_in_Block_centered + (1 + Trial_in_Block_centered *|* ID). See Appendix B for distributions of the ratings (Figure S6 and S7).

The CLMM for the trust rate detailed in Table 5 and Table 6 indicated that a two-way interaction emerged between predictability and the centered within-block trials (Estimate = 0.092, SE = 0.024, *z* = 3.776, *p < <*0.001), thus suggesting that trust changed across trials depending on whether the mobility’s motion was predictable or unpredictable. No other two-way interactions (predictability × smoothness: *p* = 0.066, smoothness × centered within-block trials: *p* = 0.651) or three-way interactions (predictability × smoothness × centered within-block trials: *p* = 0.266) reached the level of significance.

**Table 5:**
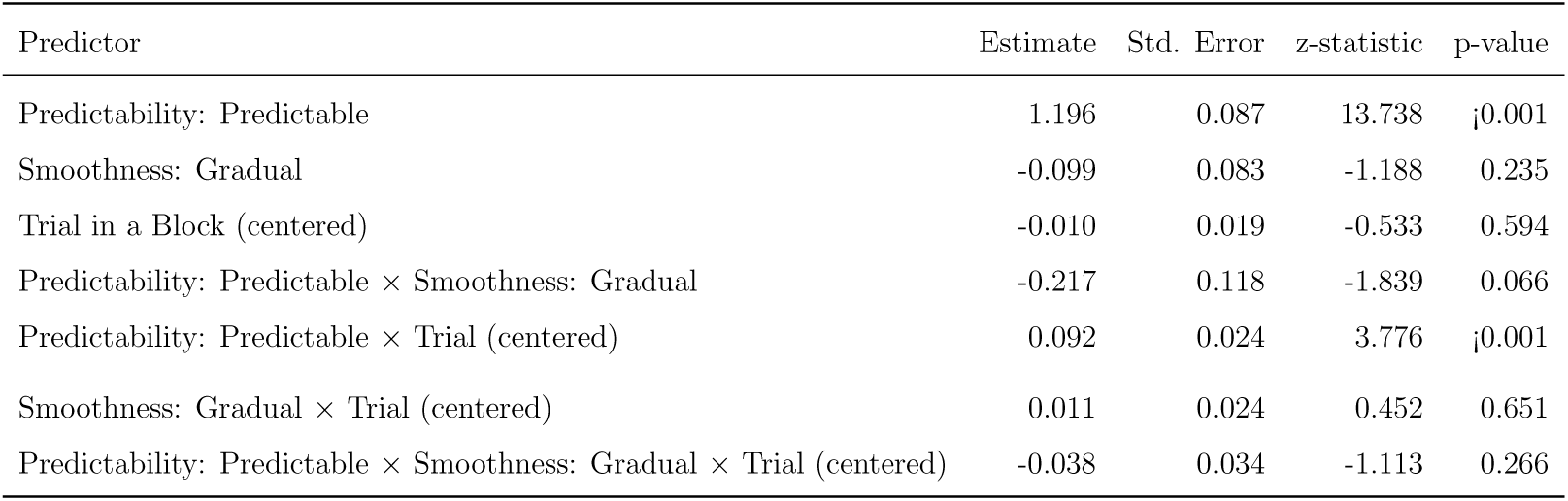
Fixed-effects coefficients of the CLMM for trust rating. The model includes predictability, smoothness, and the centered within-block trials as fixed effects, alongside ID and centered within-block trials as a random effect.

**Table 6:** Threshold estimates from the CLMM for trust rating.

| Threshold | Estimate | Std. Error | z-statistic | p-value |
| --- | --- | --- | --- | --- |
| 1—2 | -1.126 | 0.118 | -9.511 | ¡0.001 |
| 2—3 | -0.510 | 0.116 | -4.405 | ¡0.001 |
| 3—4 | 0.085 | 0.115 | 0.737 | 0.461 |
| 4—5 | 0.649 | 0.116 | 5.591 | ¡0.001 |
| 5—6 | 1.315 | 0.118 | 11.137 | ¡0.001 |
| 6—7 | 2.094 | 0.124 | 16.900 | ¡0.001 |

A simple slope analysis was conducted. This analysis estimated the trend of centered within-block trials for each level of predictability, averaging across smoothness conditions as a result of the nonsignificant higher-order interactions involving smoothness. The results are presented in Table 7. The analysis revealed that when evasions were predictable, the trust rate increased significantly as the trials progressed (slope = 0.068, SE = 0.015, *z* = 4.505, *p <*0.001). In contrast, for unpredictable evasions, the change in the trust rate over trials was not significantly different from zero (slopes = -0.005, SE = 0.015, *z* = -0.324, *p* = 0.746). The difference between these two slopes was statistically significant (*p* = 0, Tukey adjusted). Figure 7 illustrates this significant interaction, revealing a positive trend for the trust rating in the predictable condition. Note that the y-axis indicates the model’s latent scale (rather than raw trust scores).

**Figure 7:**
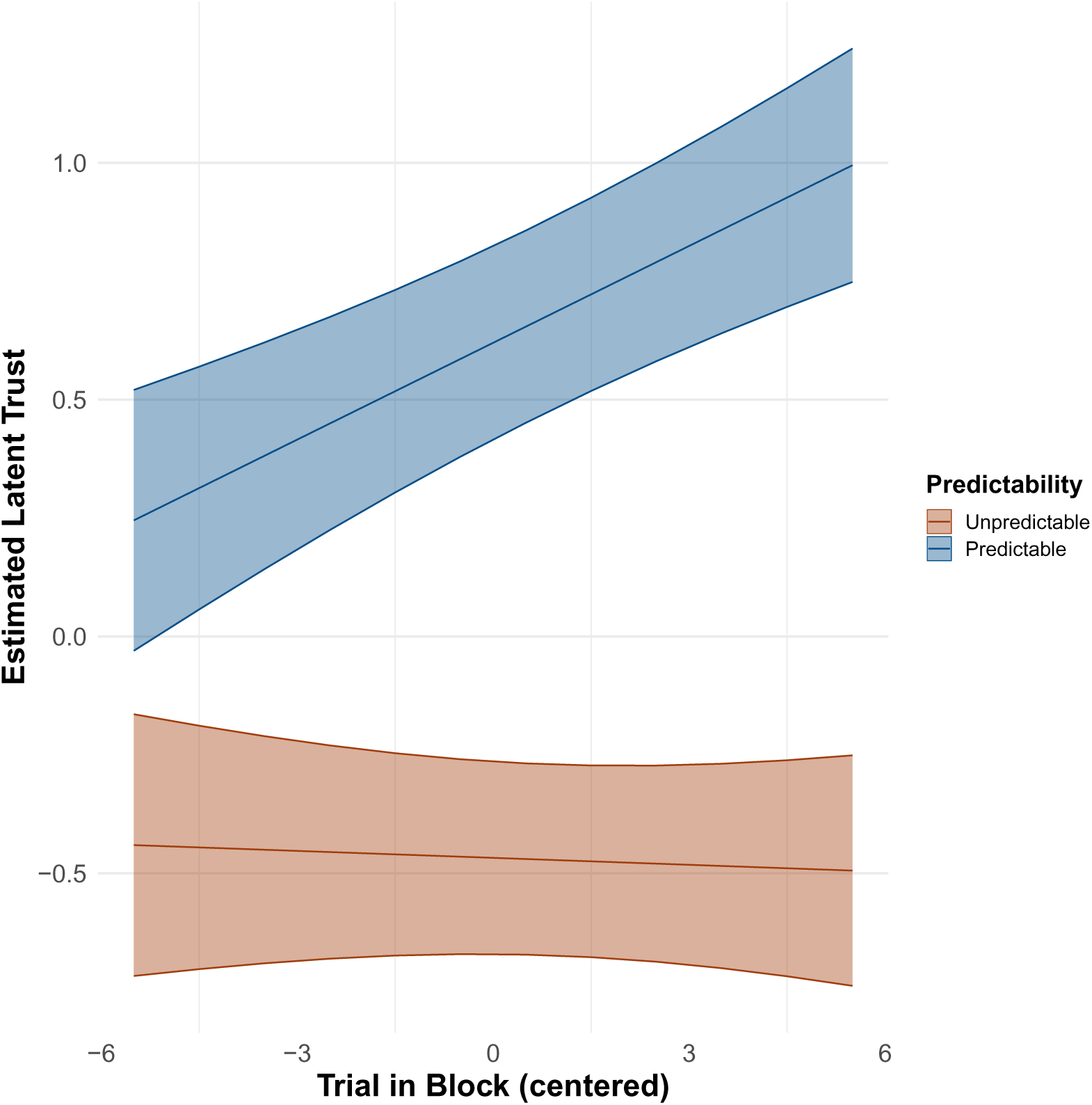
Interaction plot of the best model for the trust rate. The x-axis indicates the centered within-block trials, in which context -5.5 represents the first trial and 5.5 represents the last trial in a block. The y-axis indicates the estimated marginal means on the latent scale (linear predictor) obtained from the CLMM. The red line and area represent unpredictable conditions, whereas the blue line and area represent predictable conditions. The shaded area represents the 95% confidence intervals.

**Table 7:**
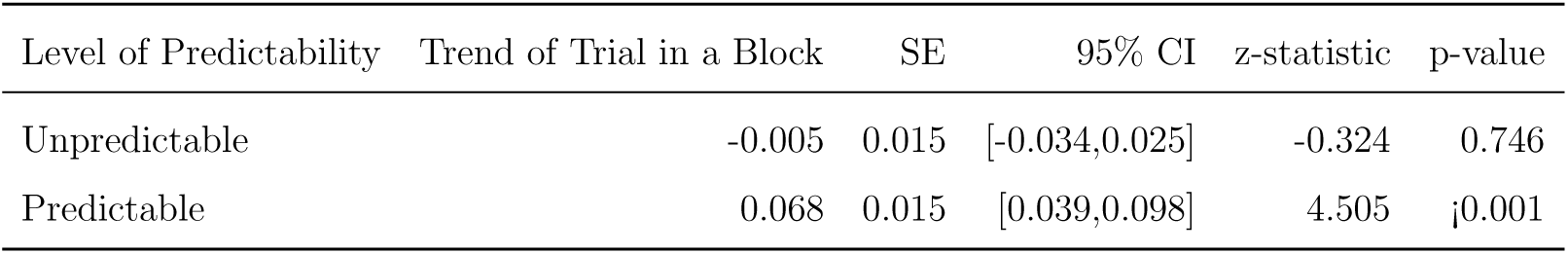
Simple slopes of the centered within-block trials for each of the four conditions.

### 5.2. Skin Conductance Responses (SCR)

A LMM was fitted to the SCR data. The final model structure was as follows: SCR ∼ Predictability * Smoothness * Trial_in_Block_centered + (1 *|* ID). However, the diagnostic checks presented in Figure S16 indicated substantial violations of the assumptions. See Appendix B for distributions of the peak and time series data (distributions in Figure S9 and S10, time series data in S8) and Appendix C for diagnostic checks (Figure S16).

To address the violations, we fitted a generalized linear mixed model (GLMM) with the assistance of the `glmer` function (provided by lme4 v1.1.35.5; (Bates et al., 2015)) with a Gamma distribution and a log link function. This choice was based on the alignment between the properties of SCR. amplitude and the Gamma distribution. The peak of the SCR amplitude is continuous, exclusively positive, and forms a positively skewed distribution. The Gamma distribution which is, a continuous probability distribution defined on positive real numbers that can represent a distribution with a long right tail, is well suited for the distribution family for such skewed data. See Appendix B for log-transformed distributions of the peak (Figure S11).

The final model structure was as follows: SCRamplitude ∼ Predictability * Smoothness * Trial_in_Block_centered + (1 + Trial_in_Block_centered *|* ID). Unlike in the case of the LMM fitted previously, diagnostic checks indicated no substantial violations of the relevant assumptions. See Appendix C for diagnostic checks (Figure S17).

Table 8 summarizes the fixed effects. The GLMM for the SCR amplitude indicated significant main effects of predictability (Estimate = -0.449, SE = 0.069, *z* = -6.469, *p <*0.001) and smoothness (Estimate = 0.142, SE = 0.069, *z* = 2.046, *p* = 0.041). Figure 8 illustrates the estimated peak of the SCR amplitude with respect to each level of predictability and smoothness, averaged across centered within-block trials. The results indicated that the SCR amplitude was significantly lower in the predictable condition than in the unpredictable condition and that it was higher in the gradual condition than in the abrupt condition. No two-way interactions (predictability × smoothness: *p* = 0.843, predictability × centered within-block trials: *p* = 0.304, smoothness × centered within-block trials: *p* = 0.717) or the three-way interaction (predictability × smoothness × centered within-block trials: *p* = 0.678) reached the level of significance.

**Figure 8:**
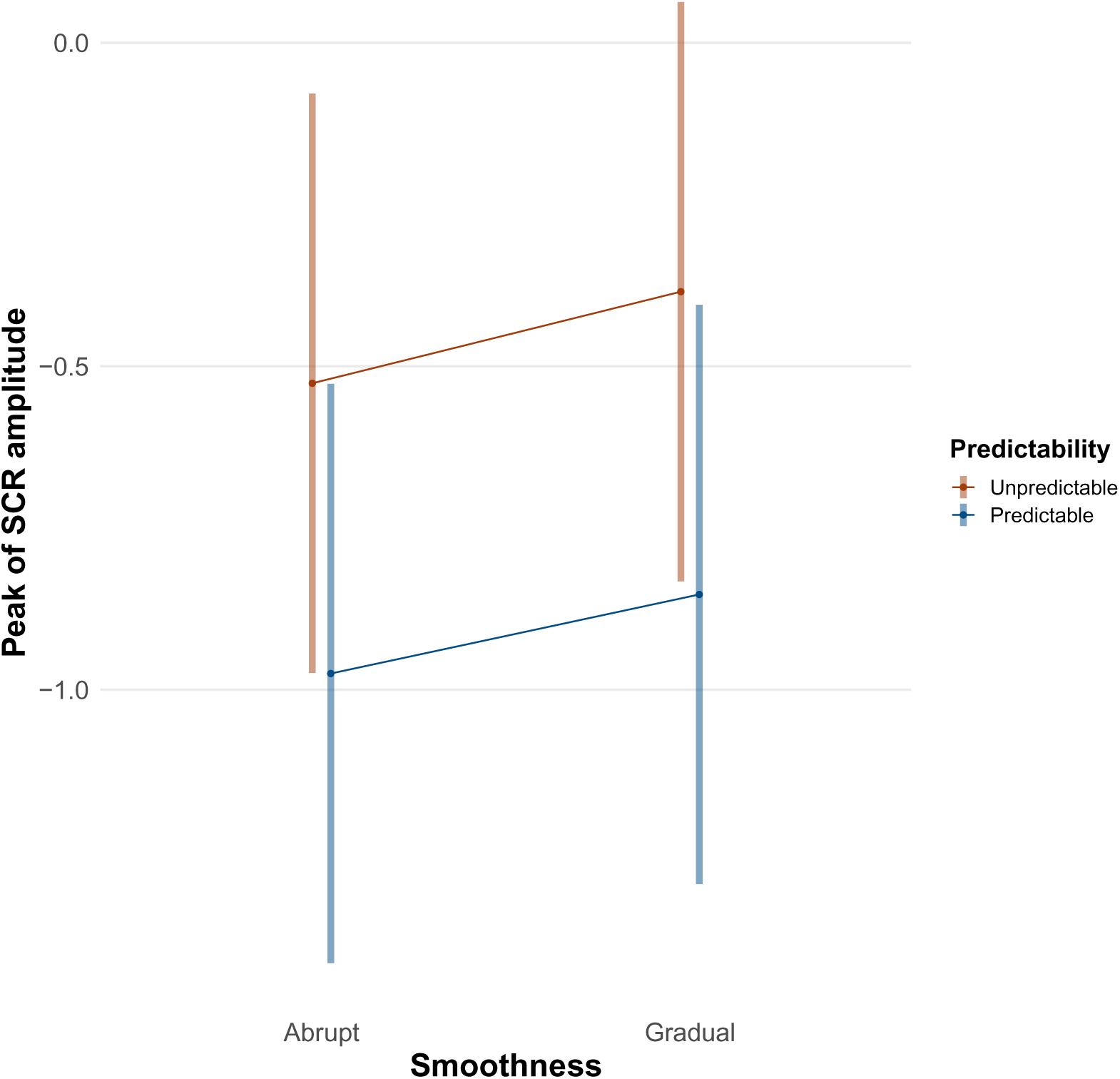
Main effects plot of the best model for SCR. The y-axis indicates the estimated peak of the SCR amplitude on its log-scale. The x-axis indicates the level of the smoothness condition. The red line and error bar represent unpredictable conditions, whereas the blue line and error bar represent Predictable conditions. The plot presents the estimated SCR amplitude as a function of predictability and smoothness. The error bars represent the 95% confidence intervals.

**Table 8:**
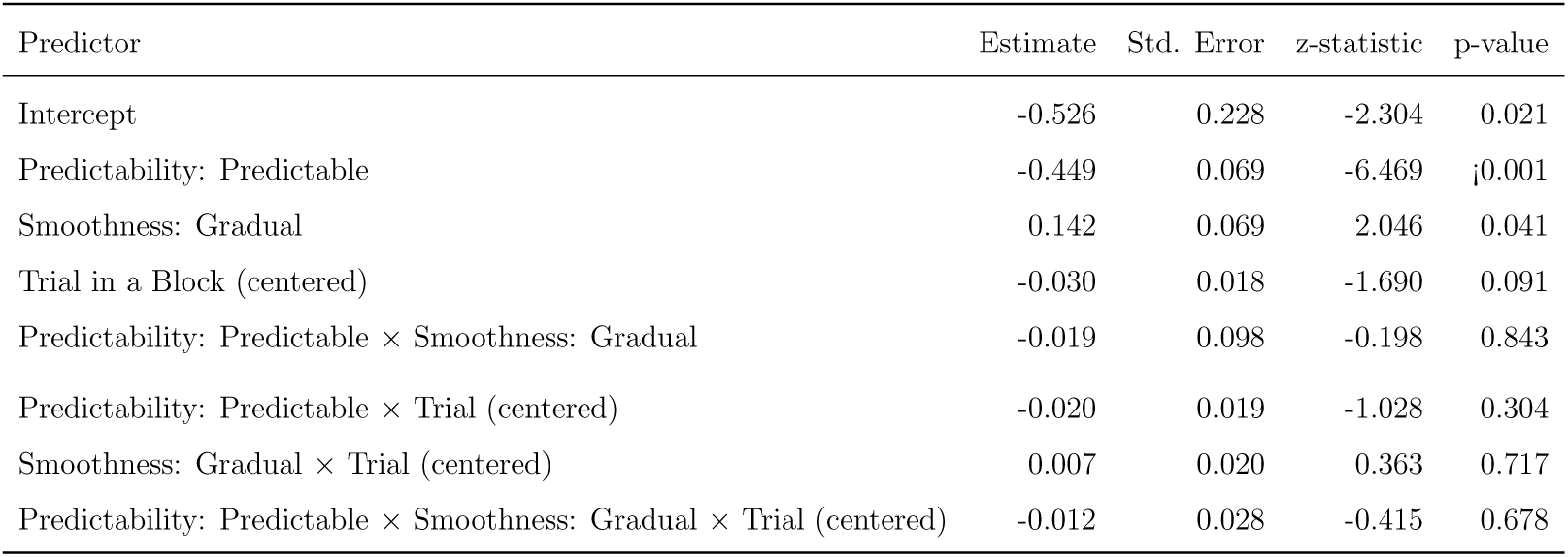
Fixed-effects coefficients of the GLMM for the peak of SCR. The model includes predictability, smoothness, and the centered within-block trials as fixed effects, alongside ID and the centered within-block trials as a random effect.

The correlation analysis between self-reported arousal and the peak of SCR is presented in in Appendix B (Figure S12 and S13).

## 6. Discussion

### 6.1. Overview of the Discussion

This study reveals that the AMR’s behavioral predictability is substantially a primary influence on users’ emotional experiences, physiological arousal, and development of trust during repeated interactions. Predictable AMR actions led to a trial-by-trial increase in positive valence. This predictability also facilitated habituation to the AMR’s movements, even when those movements were abrupt, thus leading to reduced arousal. Furthermore, predictable behavior contributed to the growth of trust over successive trials. These results are in line with research that has highlighted the importance of predictable robot movements with regard to user experience and interaction performance (Ivanova et al., 2020; Akalin et al., 2022b; Haney and Liang, 2024; Rubagotti et al., 2022).

Conversely, unpredictable AMR behavior, especially in combination with abrupt movements, increased arousal and suggested sensitization. This interaction pattern also resulted in lower levels of trust, which remained stagnant. Movement smoothness appeared to be a secondary factor; namely, it primarily influenced arousal levels when the AMR’s behavior was unpredictable, but it did not have a significant direct effect on changes in trust when predictability was taken into account.

These outcomes indicate that predictability is a dominant characteristic in the process of shaping the human-AMR relationship. These results suggest that the cognitive benefit of being able to predict the AMR’s behaviors can outweigh the startling effect of an abrupt movement. Predictability may therefore improve interactions by decreasing the cognitive load associated with uncertainty. Therefore, in terms of the implications of these findings, we suggest that a fixed avoidance rule should be preserved for the behavior of the AMR to ensure that users can learn; such a rule can include turning to the same side and a stable trigger distance. Furthermore, smoothness can be used to limit arousal when this rule cannot be applied.

### 6.2. Valence

The trial-by-trial increase in valence under predictable conditions indicated in Figure 5 and Table 2 suggests that predictability fosters positive emotional experiences via repeated interactions. These results are in line with previous findings indicating that predictable AMR behavior leads to improved performance and that robots that exhibit such behavior are even preferable to humans (Ivanova et al., 2020; Akalin et al., 2022b; Haney and Liang, 2024; Rubagotti et al., 2022). When an AMR’s actions are predictable, users can form accurate expectations of the robot, and behavior that meets these expectations can contribute to positive emotional change (Karbouj et al., 2024).

Notably, the increase in valence as trials progressed indicates an adaptive or learning process rather than a simple static preference for predictability(Churamani et al., 2022). This process represents a form of positive habituation or learning, an understanding that the interaction is safe and comfortable, which in turn enhances positive feelings over time.

While the mere-exposure effect typically refers to the ability of repeated contact to increase liking via familiarity(Bornstein and D’agostino, 1992), the results of this study suggest that, in the context of AMRs, this effect is specifically related to the predictability of the interaction. Zlotowski et al. (2015) demonstrated that AMR’s attitude, an important aspect of the nature of the interaction, significantly influences “likeability” during repeated encounters. The key attitude that enhanced “likeability”, i.e., increased valence, over time in our experiment is likely to have been the AMR’s consistent predictability.

Indeed, unpredictable encounters did not lead to similar benefits. This finding implies that mere exposure is insufficient and that the “nature” of the contact, i.e., its predictability, is critical with regard to positive emotional change.

### 6.3. Arousal

As illustrated in Figure 6, the increase in arousal that occurred as trials progressed in the unpredictable and abrupt condition indicates that this combination was particularly agitating. Unpredictable events can be perceived as potentially threatening, thus leading to higher levels of physiological and self-reported arousal (Bandura, 1988; Jackson et al., 2015). The main effect of predictability on the peak of the SCR amplitude also supports this point.

Abrupt movements can exacerbate this effect by inducing surprise (Kirschner et al., 2022). Repeated exposure to a stimulus that an individual learns is harmless leads to habituation; however, the combination of unpredictability with abruptness acts as a cumulative and notable stressor. Persistently negative and uncertain interactions that are rooted in these dual traits should lead to sensitization instead of habituation (Kirschner et al., 2021; Çevik, 2014).

In this context, users are uncertain not only regarding what would happen but also as a result of their jarring experience when the event in question does happen, thus leading to increased arousal over repeated exposures.

In contrast, the significant decrease observed in self-reported arousal across trials for both predictable conditions, i.e., predictable-abrupt and predictable-gradual, suggests the occurrence of habituation in this context. When AMR’s behavior was predictable, participants learned to anticipate its actions, even when those actions were abrupt. This learned expectation should reduce the novelty and startling effect of the movement, thereby decreasing both cognitive load and arousal(Bandura, 1988). In line with the experimental results reported by Pace-Schott et al. (2011) and Adam et al. (2015), this finding can be interpreted as indicating a decrease in arousal resulting from the repeated presentation of similar stimuli. Predictability thus facilitates habituation, thereby making learning and adaptation possible. This finding implies that the cognitive benefit of knowing what to expect can override the potentially startling effect of an abrupt movement.

Under unpredictable conditions, however, smoothness appears to modulate arousal. As indicated in the trends associated with each trial presented in Table 4, while unpredictable and abrupt movements led to an increase in arousal, unpredictable yet smooth movements did not lead to a significant change. This finding indicates that smooth motion can mitigate the arousing effects of unpredictability. A smooth but unpredictable movement may be less startling and easier to adapt to than is a movement that is both abrupt and unpredictable. The characteristics of the AMRs’ behavior under unpredictable movements, namely, in this study, its smoothness, therefore play a key role in the process of determining how arousing that movement is perceived to be.

### 6.4. Trust

The interaction between predictability and trials reported in Figure 7 and Table 7 indicates that trust is not static but rather evolves with experience. When the AMR was predictable, trust increased more substantially across trials, thus suggesting that each consistent action reinforces a user’s positive assessment of the AMR’s reliability. In contrast, under unpredictable conditions, trust began at a lower point and stagnated, as the AMR’s inconsistent actions could prevent users from confirming whether they could trust the AMR. This result is in line with findings indicating that recent failures can decrease trust (Akalin et al., 2022a). Roesler et al. (2020) also reported that a full recovery of trustworthiness becomes difficult, particularly following multiple errors. Although the unpredictable condition in this study was not designed as a “failure action”, its lack of consistency may have resulted in unpredictable movement, and such movements hindered positive adjustments in trust(Adam et al., 2015). The experimental results therefore reveal that trust formation is a dynamic process that is strongly governed by predictability; namely, predictable interactions actively build trust over time.

The interactions between smoothness and the other factors were not significant. These findings might suggest that the effect of smoothness on changes in trust is simply weak and easily masked by other influences. Previous findings indicating that performance is a more critical factor than is appearance in the context of promoting human trust in robots may explain the dominance of predictability (Hancock et al., 2011; Law et al., 2022). For example, Nordheim et al. (2019) and Law et al. (2022) reported that in the context of chatbots, performance factors such as expertise and responsiveness with accuracy are more dominant than are appearance-related factors such as human-likeness. Thus, in this study, predictability was likely perceived as a performance factor, whereas smoothness may have been viewed as more of an aesthetic element than a component of performance. Consequently, the strong effect of predictability, as the dominant factor, may have obscured the more subtle influence of smoothness on trust.

### 6.5. Skin Conductance Response (SCR)

Notably, the effect of smoothness differed between self-reported arousal and SCR. As indicated in Table 3 and Table 8, gradual motion contributed negatively to subjective arousal (i.e., a calming effect) but positively to SCR (i.e., an agitating effect). This discrepancy may be explained from the perspective of proxemics proposed by Hall (1963). The results reported by Candini et al. (2021) revealed that the amplitude of SCR increases when others approach a participant’s personal space. In our experiment, the robot in the condition involving gradual motion executed its avoidance maneuver over a longer duration, thereby maintaining a relatively small separation distance. This situation involves a deeper or more prolonged intrusion into the participant’s intimate or personal space. In the condition involving abrupt motion, conversely, the robot quickly created a large separation distance, thus making the intrusion momentary. Thus, the higher SCR associated with the gradual AMR motion could be interpreted as a physiological response to this proximity.

The divergence between these two measures may be due to the fact that subjective arousal reflects a different cognitive interpretation from the SCR. Even at close range, a gradual movement can convey a ‘nonaggressive’ or ‘controlled’ intent. For example, Hugues et al. (2016) reported that the smooth and human-like movement of a robot arm is perceived as nonaggressive, thus leading participants to feel safer and to experience less stress. Consequently, although the physiological arousal signal (the SCR investigated in this study) was generated by an intrusion on individuals’ personal space, the brain may not directly interpret that signal as indicating a state of high awareness. That is, the SCR likely responded to the direct factor of physical proximity, whereas the subjective evaluation reflected an interpretation of the movement’s low threat level, thus causing the two measures to diverge.

## 7. Limitations

This study has several limitations. First, the experimental setup entails certain constraints. The focus of this study was limited to the specific task of collision avoidance. The results might vary in other contexts involving HRI, such as collaborative assembly or social companionship. Furthermore, the morphology of AMRss may have influenced the corresponding outcomes, as responses can change alongside different forms of AMR, such as humanoid robots or animal-like models.

Second, the measurement methods used in this research entail certain limitations. Valence and arousal were measured via self-reports, a common method that is nevertheless susceptible to bias. The low marginal *R*^2^ (0.022) and higher conditional *R*^2^ (0.439) suggest substantial individual variation in the relationship between perceived arousal and peak SCR amplitude, as illustrated in S12. Factors such as individual differences in emotional recognition, personal interpretations of the EmojiGrid(Toet et al., 2018, 2019), or other unmeasured cognitive assessments may also have influenced these self-reports.

Third, the study was limited in terms of its ability to investigate longterm effects. The experiment involved short-term interactions on the basis of 48 trials per participant. The effects of long-term interactions might differ, as familiarity and adaptation could evolve over longer periods.

## 8. Conclusion

This study demonstrated that predictability is a consistently important and dominant factor in terms of valence, arousal, and trust. Predictable AMR behavior fostered more positive changes in valence, decreased arousal through habituation, and increased the growth of trust as trials progressed. This consistency suggests that designing for predictability is a robust strategy that can be used to improve the user experience in the context of HRI.

Furthermore, abrupt motion, particularly during unpredictable behavior, led to sensitization rather than habituation over time. This finding indicates that motion smoothness plays a moderating role in the extent to which humans perceive unpredictable movements as arousing.

The findings of this study suggest that designers and engineers should prioritize the task of ensuring that robot behavior is predictable or can be learned quickly. While smoothness can modulate arousal in unpredictable scenarios, its effects on valence and trust were less pronounced. We should therefore focus on the task of ensuring that robots’ intentions and future actions are communicated clearly and that of ensuring that their behavior is inherently consistent. This approach can allow users to form accurate expectations, which in turn should foster positive emotions and trust in the context of their interactions with the robot.

## Statements and Declarations

### Acknowledgments

This work was supported by JSPS KAKENHI (Grant Numbers JP25K21323 to H.T., JP25H01141 to S.N., and JP23KK0183 to T.M.), the Nitto Foundation, and the Amano Institute of Technology.

### Data statement

The code and data underlying the results presented in the study are available from the Open Science Framework repository (https://osf.io/tr85q/).

### Artificial intelligence

During the preparation of this work, the authors used Gemini 2.5 Pro and Grammarly to improve the language. The authors reviewed and edited the content as needed and take full responsibility for the content of the publication.

### Authorship contribution statement

**Yuta Matsubara:** Conceptualization, data curation, formal analysis, investigation, methodology, software, visualization, writing - original draft. **Hideki Tamura:** Conceptualization, funding acquisition, methodology, project administration, resources, supervision, validation, writing - review & editing. **Tetsuto Minami:** Funding acquisition, project administration, resources, supervision, validation, writing - review & editing. **Shigeki Nakauchi:** Funding acquisition, project administration, resources, supervision, validation, writing - review & editing

HRI: human-robot interaction
AMR: autonomous mobile robot
SCR: skin conductance response
LMM: linear mixed model
GLMM: generalized linear mixed model
CLMM: cumulative link mixed model

## References

Adam, M. T. P., Krämer, J., and Müller, M. B. (2015). Auction Fever! How Time Pressure and Social Competition Affect Bidders ‘Arousal and Bids in Retail Auctions. Journal of Retailing, 91(3):468–485. Publisher: JAI.

Akaike, H. (1987). Factor analysis and AIC. Psychometrika, 52(3):317–332.

Akalin, N., Kiselev, A., Kristoffersson, A., and Loutfi, A. (2023). A Taxonomy of Factors Influencing Perceived Safety in Human–Robot Interaction. International Journal of Social Robotics, 15(12):1993–2004.

Akalin, N., Kristoffersson, A., and Loutfi, A. (2022a). Do you feel safe with your robot? Factors influencing perceived safety in human-robot interaction based on subjective and objective measures. International Journal of Human-Computer Studies, 158:102744.

Akalin, N., Kristoffersson, A., and Loutfi, A. (2022b). Do you feel safe with your robot? Factors influencing perceived safety in human-robot interaction based on subjective and objective measures. International Journal of Human-Computer Studies, 158:102744.

Akinade, A., Barros, D., and Vernon, D. (2025). Biological Motion Aids Gestural Communication by Humanoid Social Robots. International Journal of Humanoid Robotics, 22(02):2550001.

Alhaji, B., Büttner, S., Sanjay Kumar, S., and Prilla, M. (2025). Trust dynamics in human interaction with an industrial robot. Behaviour & Information Technology, 44(2):266–288.

Apraiz, A., Lasa, G., Mazmela, M., Arana-Arexolaleiba, N., Serrano Muñoz, A., Elguea, c., and Etxabe, A. (2025). Evaluating the Effect of Speed and Acceleration on Human Factors during an Assembly Task in Human–Robot Interaction (HRI). International Journal of Social Robotics, 17(2):211–256.

Ashraf, N., Bohnet, I., and Piankov †, N. (2006). Decomposing Trust and Trustworthiness. Experimental Economics, 9(3):193–208.

Babel, F., Kraus, J., and Baumann, M. (2022). Findings from a qualitative field study with an autonomous robot in public: Exploration of user reactions and conflicts. International Journal of Social Robotics, 14(7):1625–1655.

Bandura, A. (1988). Self-efficacy conception of anxiety. Anxiety Research, 1(2):77–98. Publisher: Routledge eprint: 10.1080/10615808808248222.

Bartneck, C., Kulíc, D., Croft, E., and Zoghbi, S. (2009). Measurement Instruments for the Anthropomorphism, Animacy, Likeability, Perceived Intelligence, and Perceived Safety of Robots. International Journal of Social Robotics, 1(1):71–81.

Bates, D., Mächler, M., Bolker, B., and Walker, S. (2015). Fitting linear mixed-effects models using lme4. Journal of Statistical Software, 67(1):1–48.

Berton, F., Olivier, A. H., Bruneau, J., Hoyet, L., and Pettre, J. (2019). Studying gaze behaviour during collision avoidance with a virtual walker: Influence of the virtual reality setup. 26th IEEE Conference on Virtual Reality and 3D User Interfaces, VR 2019 - Proceedings, pages 717–725. ISBN: 9781728113777 Publisher: Institute of Electrical and Electronics Engineers Inc.

Bethel, C. L., Salomon, K., Murphy, R. R., and Burke, J. L. (2007). Survey of psychophysiology measurements applied to human-robot interaction. Proceedings - IEEE International Workshop on Robot and Human Interactive Communication, pages 732–737. ISBN: 1424416345.

Bornstein, R. F. and D’agostino, P. R. (1992). Stimulus recognition and the mere exposure effect. Journal of personality and social psychology, 63(4):545.

Bortot, D., Born, M., and Bengler, K. (2013). Directly or on detours? How should industrial robots approximate humans? In 2013 8th ACM/IEEE International Conference on Human-Robot Interaction (HRI), pages 89–90. ISSN: 2167-2148.

Boucsein, W. (2012). Principles of Electrodermal Phenomena. In Boucsein, W., editor, Electrodermal Activity, pages 1–86. Springer US, Boston, MA.

Bradley, M. M. and Lang, P. J. (1994). Measuring emotion: The self-assessment manikin and the semantic differential. Journal of Behavior Therapy and Experimental Psychiatry, 25(1):49–59.

Braithwaite, J. J., Watson, D. G., Jones, R., and Rowe, M. (2013). A guide for analysing electrodermal activity (EDA) & skin conductance responses (SCRs) for psychological experiments. Psychophysiology, 49(1):1017–1034.

Camara, F. and Fox, C. (2022). Unfreezing autonomous vehicles with game theory, proxemics, and trust. Frontiers in Computer Science, 4. Publisher: Frontiers.

Campagna, G. and Rehm, M. (2024). A Systematic Review of Trust Assessments in Human-Robot Interaction. J. Hum.-Robot Interact. Just Accepted.

Candini, M., Battaglia, S., Benassi, M., di Pellegrino, G., and Frassinetti, F. (2021). The physiological correlates of interpersonal space. Scientific Reports, 11(1):2611. Publisher: Nature Publishing Group.

Çevik, M. Ö. (2014). Habituation, sensitization, and Pavlovian conditioning. Frontiers in Integrative Neuroscience, 8:13.

Chen, N., Mohanty, S., Jiao, J., and Fan, X. (2021). To err is human: Tolerate humans instead of machines in service failure. Journal of Retailing and Consumer Services, 59:102363.

Christensen, R. H. B. (2023). ordinal—Regression Models for Ordinal Data. R package version 2023.12–4.1.

Churamani, N., Barros, P., Gunes, H., and Wermter, S. (2022). Affect-Driven Learning of Robot Behaviour for Collaborative Human-Robot Interactions. Frontiers in Robotics and AI, 9. Publisher: Frontiers.

Daronnat, S., Azzopardi, L., Halvey, M., and Dubiel, M. (2021). Inferring Trust From Users ‘Behaviours; Agents ‘Predictability Positively Affects Trust, Task Performance and Cognitive Load in Human-Agent Real-Time Collaboration. Frontiers in Robotics and AI, 8. Publisher: Frontiers.

Dragan, A. and Srinivasa, S. (2013). Generating legible motion.

Duraklı, Z. and Nabiyev, V. (2022). A new approach based on Bezier curves to solve path planning problems for mobile robots. Journal of Computational Science, 58:101540.

Dzindolet, M. T., Peterson, S. A., Pomranky, R. A., Pierce, L. G., and Beck, H. P. (2003). The role of trust in automation reliance. International Journal of Human-Computer Studies, 58(6):697–718.

Firmino de Souza, D., Sousa, S., Kristjuhan-Ling, K., Dunajeva, O., Roosileht, M., Pentel, A., Mõttus, M., Can Özdemir, M., and Graťsjova, c. (2025). Trust and Trustworthiness from Human-Centered Perspective in Human–Robot Interaction (HRI)―A Systematic Literature Review. Electronics, 14(8):1557. Number: 8 Publisher: Multidisciplinary Digital Publishing Institute.

Greenberg, B., Gonźalez-Bravo, U., Yi, J., and Feldman, J. (2025). Effects of Mobile Robot Passing-Motion Path Curvature on Human Affective States in a Hallway Environment. International Journal of Social Robotics.

Guillén Ruiz, S., Calderita, L. V., Hidalgo-Paniagua, A., and Bandera Rubio, J. P. (2020). Measuring Smoothness as a Factor for Efficient and Socially Accepted Robot Motion. Sensors, 20(23):6822. Number: 23 Publisher: Multidisciplinary Digital Publishing Institute.

Gupta, R., Shin, H., Norman, E., Stephens, K. K., Lu, N., and Sentis, L. (2024). Human Stress Response and Perceived Safety during Encounters with Quadruped Robots. In 2024 33rd IEEE International Conference on Robot and Human Interactive Communication (ROMAN), pages 783–790. ISSN: 1944-9437.

Hall, E. T. (1963). A System for the Notation of Proxemic Behavior. American Anthropologist, 65(5):1003–1026. eprint: https://onlinelibrary.wiley.com/doi/pdf/10.1525/aa.1963.65.5.02a00020.

Hancock, P. A., Billings, D. R., Schaefer, K. E., Chen, J. Y. C., de Visser, E. J., and Parasuraman, R. (2011). A Meta-Analysis of Factors Affecting Trust in Human-Robot Interaction. Human Factors, 53(5):517–527. Publisher: SAGE Publications Inc.

Haney, J. M. and Liang, C.-J. (2024). A Literature Review on Safety Perception and Trust during Human–Robot Interaction with Autonomous Mobile Robots That Apply to Industrial Environments. IISE Transactions on Occupational Ergonomics and Human Factors, 12(1-2):6–27. Publisher: Taylor & Francis eprint: 10.1080/24725838.2023.2283537.

Hugues, O., Weistroffer, V., Paljic, A., Fuchs, P., Karim, A. A., Gaudin, T., and Buendia, A. (2016). Determining the Important Subjective Criteria in the Perception of Human-Like Robot Movements Using Virtual Reality. International Journal of Humanoid Robotics, 13(02):1550033. Publisher: World Scientific Publishing Co.

Ivanova, E., Carboni, G., Eden, J., Krüger, J., and Burdet, E. (2020). For Motion Assistance Humans Prefer to Rely on a Robot Rather Than on an Unpredictable Human. IEEE Open Journal of Engineering in Medicine and Biology, 1:133–139.

Jackson, F., Nelson, B. D., and Proudfit, G. H. (2015). In an uncertain world, errors are more aversive: Evidence from the error-related negativity. Emotion, 15(1):12–16. Place: US Publisher: American Psychological Association.

Juuse, L., Tamm, D., Lõo, K., Allik, J., and Kreegipuu, K. (2024). Skin conductance response and habituation to emotional facial expressions and words. Acta Psychologica, 251:104573.

Kandul, S., Micheli, V., Beck, J., Burri, T., Fleuret, F., Kneer, M., and Christen, M. (2023). Human control redressed: Comparing AI and human predictability in a real-effort task. Computers in Human Behavior Reports, 10:100290.

Karbouj, B., Alshamaa, O., Al Rashwany, K., and Krüger, J. (2024). Enhancing Human-Robot Collaborative Predictability through Rational Action Modeling of Robot Trajectories. Procedia CIRP, 130:516–523.

Kirschner, R. J., Burr, L., Porzenheim, M., Mayer, H., Abdolshah, S., and Haddadin, S. (2021). Involuntary Motion in Human-Robot Interaction: Effect of Interactive User Training on the Occurrence of Human Startle-Surprise Motion. In 2021 IEEE International Conference on Intelligence and Safety for Robotics (ISR), pages 28–32.

Kirschner, R. J., Mayer, H., Burr, L., Mansfeld, N., Abdolshah, S., and Haddadin, S. (2022). Expectable Motion Unit: Avoiding Hazards From Human Involuntary Motions in Human-Robot Interaction. IEEE Robotics and Automation Letters, 7(2):2993–3000.

Kohn, S. C., de Visser, E. J., Wiese, E., Lee, Y.-C., and Shaw, T. H. (2021). Measurement of Trust in Automation: A Narrative Review and Reference Guide. Frontiers in Psychology, 12.

Kok, B. C. and Soh, H. (2020a). Trust in Robots: Challenges and Opportunities. Current Robotics Reports, 1(4):297–309.

Kok, B. C. and Soh, H. (2020b). Trust in Robots: Challenges and Opportunities. Current Robotics Reports, 1(4):297–309.

Kraus, J., Babel, F., Hock, P., Hauber, K., and Baumann, M. (2022). The trustworthy and acceptable HRI checklist (TA-HRI): questions and design recommendations to support a trust-worthy and acceptable design of human-robot interaction. Gruppe. Interaktion. Organisation. Zeitschrift für Angewandte Organisationspsychologie (GIO), 53(3):307–328.

Lasota, P. A., Fong, T., Shah, J. A., and Delft, B. (2017). A Survey of Methods for Safe Human-Robot Interaction. Foundations and Trends® in Robotics, 5(4):261–349. ISBN: 9781680832785 Publisher: Now Publishers, Inc.

Law, E. L.-C., FØLstad, A., and Van As, N. (2022). Effects of Humanlikeness and Conversational Breakdown on Trust in Chatbots for Customer Service. In Nordic Human-Computer Interaction Conference, NordiCHI ‘22, pages 1–13, New York, NY, USA. Association for Computing Machinery.

Lüdecke, D., Ben-Shachar, M. S., Patil, I., Waggoner, P., and Makowski, D. (2021). performance: An R package for assessment, comparison and testing of statistical models. Journal of Open Source Software, 6(60):3139.

LeDoux, J. E. and Hofmann, S. G. (2018). The subjective experience of emotion: a fearful view. Current opinion in behavioral sciences, 19:67–72.

Lee, J. D. and See, K. A. (2004). Trust in Automation: Designing for Appropriate Reliance. Human Factors, 46(1):50–80. Publisher: SAGE Publications Inc.

Lenth, R. V. (2024). emmeans: Estimated Marginal Means, aka Least-Squares Means. R package version 1.10.5.

Li, R., van Almkerk, M., van Waveren, S., Carter, E., and Leite, I. (2019). Comparing human-robot proxemics between virtual reality and the real world. In 2019 14th ACM/IEEE International Conference on Human-Robot Interaction (HRI), pages 431–439.

Li, Y., Wu, B., Huang, Y., and Luan, S. (2024). Developing trustworthy artificial intelligence: insights from research on interpersonal, human-automation, and human-AI trust. Frontiers in Psychology, 15:1382693.

Lingam, S. N., Petermeijer, S. M., Torre, I., Bazilinskyy, P., Ljungblad, S., and Martens, M. (2025). Behavioral Effects of a Delivery Drone on Feelings of Uncertainty: A Virtual Reality Experiment. J. Hum.-Robot Interact. Just Accepted.

Lonsdorf, T. B., Klingelhöfer-Jens, M., Andreatta, M., Beckers, T., Chalkia, A., Gerlicher, A., Jentsch, V. L., Drexler, S. M., Mertens, G., Richter, J., Sjouwerman, R., Wendt, J., and Merz, C. J. (2019). Navigating the garden of forking paths for data exclusions in fear conditioning research. eLife, 8. Publisher: eLife Sciences Publications Ltd.

Malle, B. F. and Ullman, D. (2021). Chapter 1–a multidimensional conception and measure of human-robot trust. Trust in Human-Robot Interaction, pages 3–25.

Mara, M. and Meyer, K. (2022). Acceptance of Autonomous Vehicles: An Overview of User-Specific, Car-Specific and Contextual Determinants. In User Experience Design in the Era of Automated Driving, pages 51–83. Springer International Publishing.

Mayer, R., Davis, J., and Schoorman, F. (1995). An Integrative Model of Organizational Trust. ACADEMY OF MANAGEMENT REVIEW, 20(3):709–734. Num Pages: 26 Place: Briarcliff Manor Publisher: Acad Management Web of Science ID: WOS:A1995RJ62200009.

Neath, A. A. and Cavanaugh, J. E. (2012). The Bayesian information criterion: background, derivation, and applications. WIREs Computational Statistics, 4(2):199–203. eprint: https://onlinelibrary.wiley.com/doi/pdf/10.1002/wics.199.

Neggers, M. M. E., Belgers, S., Cuijpers, R. H., Ruijten, P. A. M., and IJsselsteijn, W. A. (2024). Comfortable Crossing Strategies for Robots. International Journal of Social Robotics, 16(7):1541–1560.

Nicholls, M. E., Thomas, N. A., Loetscher, T., and Grimshaw, G. M. (2013). The flinders handedness survey (flanders): a brief measure of skilled hand preference. Cortex, 49(10):2914–2926.

Nordheim, C. B., Følstad, A., and Bjørkli, C. A. (2019). An Initial Model of Trust in Chatbots for Customer Service―Findings from a Questionnaire Study. Interacting with Computers, 31(3):317–335.

Núnez, P., Manso, L. J., Bustos, P., Drews Jr, P., and Macharet, D. G. (2016). Towards a new semantic social navigation paradigm for autonomous robots using cortex. In IEEE international symposium on robot and human interactive communication (RO-MAN 2016)―BAILAR2016 workshop.

Okubo, M., Suzuki, H., and Nicholls, M. (2014). A japanese version of the flanders handedness questionnaire. Shinrigaku Kenkyu: The Japanese Journal of Psychology, 85(5):474–481.

Pace-Schott, E. F., Shepherd, E., Spencer, R. M. C., Marcello, M., Tucker, M., Propper, R. E., and Stickgold, R. (2011). Napping promotes intersession habituation to emotional stimuli. Neurobiology of Learning and Memory, 95(1):24–36. Publisher: Academic Press.

Pauw, L. S., Sauter, D. A., van Kleef, G. A., Lucas, G. M., Gratch, J., and Fischer, A. H. (2022). The avatar will see you now: Support from a virtual human provides socio-emotional benefits. Computers in Human Behavior, 136:107368.

Plaks, J. E., Bustos Rodriguez, L., and Ayad, R. (2022). Identifying psychological features of robots that encourage and discourage trust. Computers in Human Behavior, 134:107301.

Roesler, E., Onnasch, L., and Majer, J. I. (2020). The Effect of Anthropomorphism and Failure Comprehensibility on Human-Robot Trust. Proceedings of the Human Factors and Ergonomics Society Annual Meeting, 64(1):107–111. Publisher: SAGE Publications Inc.

Rossi, A., Dautenhahn, K., Koay, K. L., and Walters, M. L. (2017). How the Timing and Magnitude of Robot Errors Influence Peoples Trust of Robots in an Emergency Scenario. Social Robotics, pages 42–52.

Rubagotti, M., Tusseyeva, I., Baltabayeva, S., Summers, D., and Sandygulova, A. (2022). Perceived safety in physical human–robot interaction―A survey. Robotics and Autonomous Systems, 151:104047.

S. M. B. P. B., S., Valori, M., Legnani, G., and Fassi, I. (2025). Assessing Safety in Physical Human–Robot Interaction in Industrial Settings: A Systematic Review of Contact Modelling and Impact Measuring Methods. Robotics, 14(3):27. Number: 3 Publisher: Multidisciplinary Digital Publishing Institute.

Saßmannshausen, T., Burggräf, P., Hassenzahl, M., and Wagner, J. (2023). Human trust in otherware–a systematic literature review bringing all antecedents together. Ergonomics, 66(7):976–998.

Shariati, A., Shahab, M., Meghdari, A., Amoozandeh Nobaveh, A., Rafatnejad, R., and Mozafari, B. (2018). Virtual Reality Social Robot Platform: A Case Study on Arash Social Robot. In Ge, S. S., Cabibihan, J.-J., Salichs, M. A., Broadbent, E., He, H., Wagner, A. R., and Castro-González, c., editors, Social Robotics, pages 551–560, Cham. Springer International Publishing.

Simões, A. C., Pinto, A., Santos, J., Pinheiro, S., and Romero, D. (2022). Designing human-robot collaboration (HRC) workspaces in industrial settings: A systematic literature review. Journal of Manufacturing Systems, 62:28–43.

Souza, D. F. d., Sousa, S., Kristjuhan-Ling, K., Dunajeva, O., Roosileht, M., Pentel, A., Mõttus, M., Özdemir, M. C., and Gratšjova, c. (2025). Trust and Trustworthiness from Human-Centered Perspective in HRI – A Systematic Literature Review. arXiv:2501.19323 [cs].

Spezialetti, M., Placidi, G., and Rossi, S. (2020). Emotion Recognition for Human-Robot Interaction: Recent Advances and Future Perspectives. Frontiers in Robotics and AI, 7. Publisher: Frontiers.

Stower, R., Kappas, A., and Sommer, K. (2024). When is it right for a robot to be wrong? Children trust a robot over a human in a selective trust task. Computers in Human Behavior, 157:108229.

Swangnetr, M. and Kaber, D. B. (2013). Emotional state classification in patient-robot interaction using wavelet analysis and statistics-based feature selection. IEEE Transactions on Human-Machine Systems, 43(1):63–75.

Tiberio, L., Cesta, A., and Belardinelli, M. O. (2013). Psychophysiological Methods to Evaluate User ‘s Response in Human Robot Interaction: A Review and Feasibility Study. Robotics 2013, Vol. 2, Pages 92-121, 2(2):92–121. Publisher: Multidisciplinary Digital Publishing Institute.

Toet, A., Heijn, F., Brouwer, A. M., Mioch, T., and Erp, J. B. F. v. (2019). The EmojiGrid as an Immersive Self-report Tool for the Affective Assessment of 360 VR Videos. Lecture Notes in Computer Science (including subseries Lecture Notes in Artificial Intelligence and Lecture Notes in Bioinformatics), 11883 LNCS:330–335. ISBN: 9783030319076 Publisher: Springer.

Toet, A., Kaneko, D., Ushiama, S., Hoving, S., Kruijf, I. d., Brouwer, A. M., Kallen, V., and Erp, J. B. F. v. (2018). EmojiGrid: A 2D Pictorial Scale for the Assessment of Food Elicited Emotions. Frontiers in Psychology, 9(NOV). Publisher: Frontiers Media SA.

van den Brule, R., Dotsch, R., Bijlstra, G., Wigboldus, D. H. J., and Haselager, P. (2014). Do Robot Performance and Behavioral Style affect Human Trust? International Journal of Social Robotics, 6(4):519–531.

Vassallo, C., Olivier, A.-H., Souères, P., Crétual, A., Stasse, O., and Pettré, J. (2017). How do walkers avoid a mobile robot crossing their way? Gait & Posture, 51:97–103.

Vassallo, C., Olivier, A.-H., Souères, P., Crétual, A., Stasse, O., and Pettré, J. (2018). How do walkers behave when crossing the way of a mobile robot that replicates human interaction rules? Gait & Posture, 60:188–193.

Ventura-Bort, C., Wendt, J., and Weymar, M. (2022). New insights on the correspondence between subjective affective experience and physiological responses from representational similarity analysis. Psychophysiology, 59(11):e14088. Publisher: John Wiley & Sons, Ltd.

Wang, Y., Li, X., Zhang, J., Li, S., Xu, Z., and Zhou, X. (2021). Review of wheeled mobile robot collision avoidance under unknown environment. Science Progress, 104(3):00368504211037771.

Westfall, J. (2015). Pangea: Power analysis for general anova designs. Unpublished manuscript. Available at http://jakewestfall.org/publications/-pangea.pdf, 4.

Yam, K. C., Tang, P. M., Jackson, J. C., Su, R., and Gray, K. (2023). The rise of robots increases job insecurity and maladaptive workplace behaviors: Multimethod evidence. The Journal of Applied Psychology, 108(5):850–870.

Yamauchi, T., Tamura, H., Minami, T., and Nakauchi, S. (2025). Waist rotation angle as indicator of probable human collision-avoidance direction for autonomous mobile robots. PloS one, 20(5):e0323632.

Zlotowski, J. A., Sumioka, H., Nishio, S., Glas, D. F., Bartneck, C., and Ishiguro, H. (2015). Persistence of the uncanny valley: the influence of repeated interactions and a robot’s attitude on its perception. Frontiers in Psychology, 6. Publisher: Frontiers.

