## Supplementary Information for "Human trust and emotion in the context of collision avoidance with an autonomous mobile robot: An investigation of predictability and smoothness in virtual reality"

### Supplement

#### *S.1. Virtual Reality Equipment*

Figure S1 shows the head-mounted display and the controller used in the experiment.

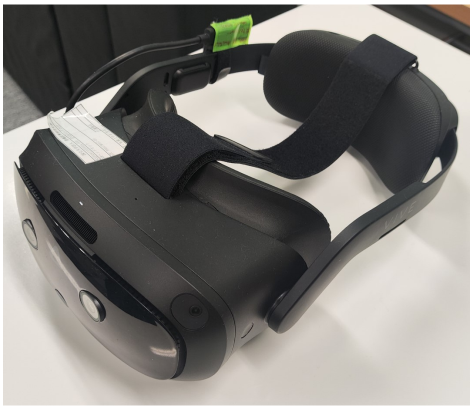

(a) The head-mounted display used in the experiment.

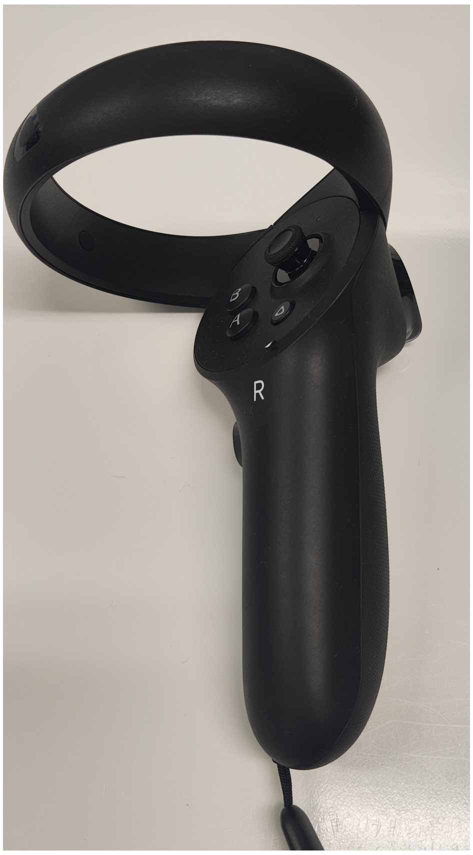


(b) The controller used by the participant to answer their emotion and trust ratings.

Figure S1: The VR equipment used in the experiment.

#### *S.2. Detailed Distribution of the Results*

##### *S.2.1. Valence*

Figure S2 and S3 depicts the distribution of valence scored by the participants for each of the four conditions and the distribution throughout the trials in each condition, respectively.

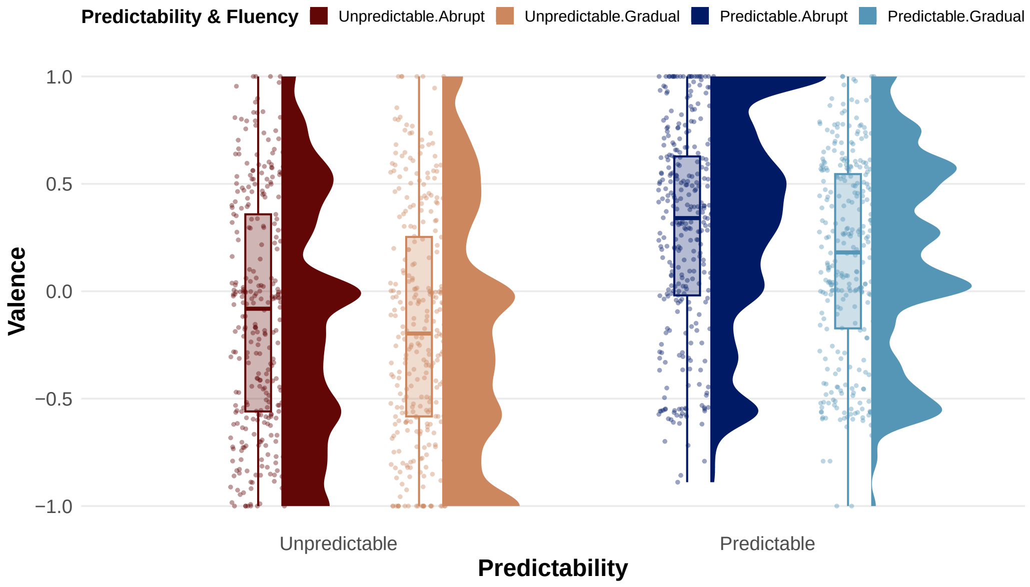


Figure S2: Raincloud plots of subjective valence ratings. The x-axis displays the combinations of each level of the conditions. The y-axis represents the valence rating on a continuous scale from -1 (most negative) to 1 (most positive).

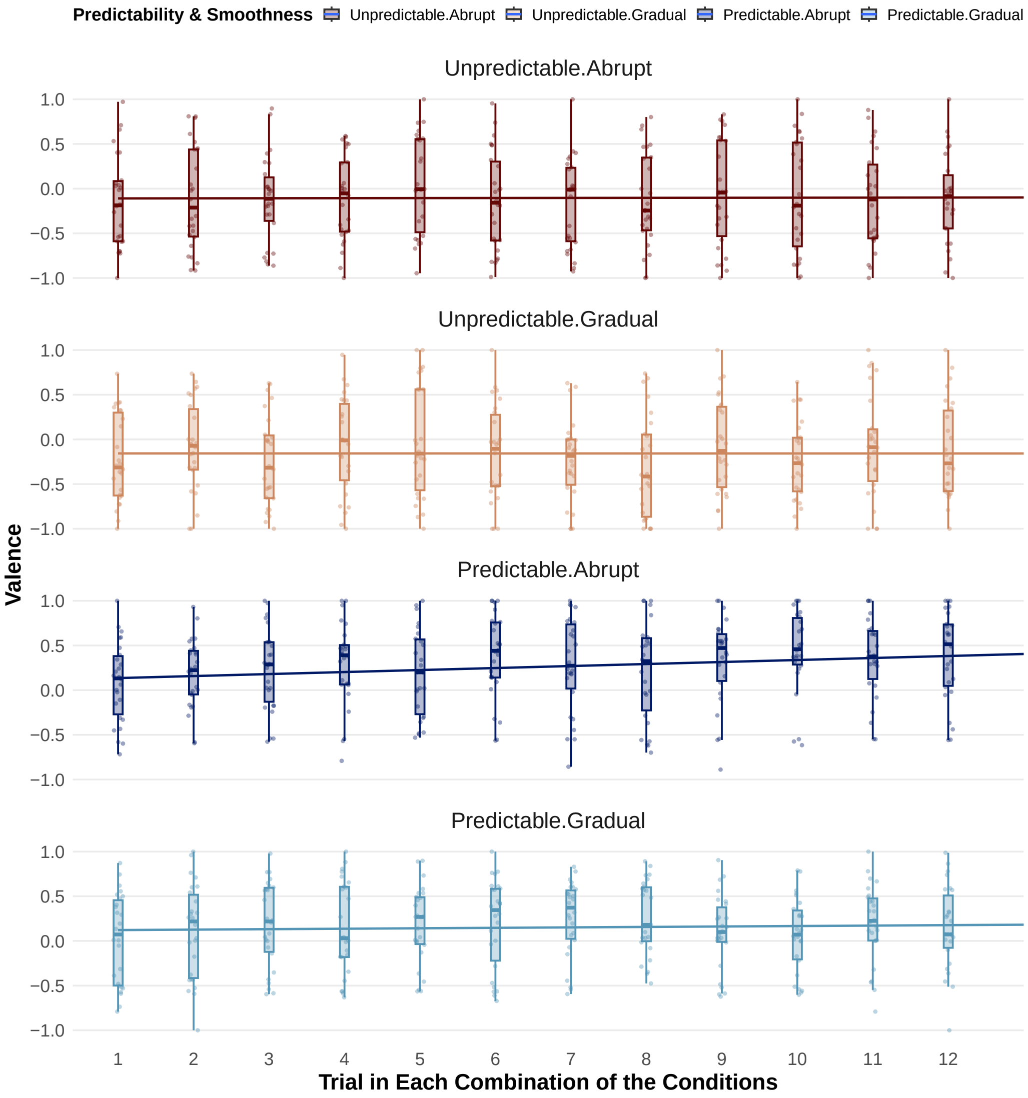


Figure S3: Scattered plots with box plots and regression lines of subjective ratings of valence in each combination of the conditions. The x-axis displays the trials in a block associated with each combination of the conditions. The y-axis represents the valence rating.

The mean and standard error of valence rating for each condition were as follows: `Unpredictable-Abrupt`: -0.104 $\pm$ 0.030, `Unpredictable-Gradual`: -0.157 $\pm$ 0.030, `Predictable-Abrupt`: 0.281 $\pm$ 0.027, `Predictable-Gradual`:

0.154 ± 0.026. For the summary for each of the four conditions throughout the trials, see Supplement D.

#### *S.2.2. Arousal*

Figure S4 and S5 depicts the distribution of arousal scored by the participants for each of the four conditions and the distribution throughout the trials in each condition, respectively.

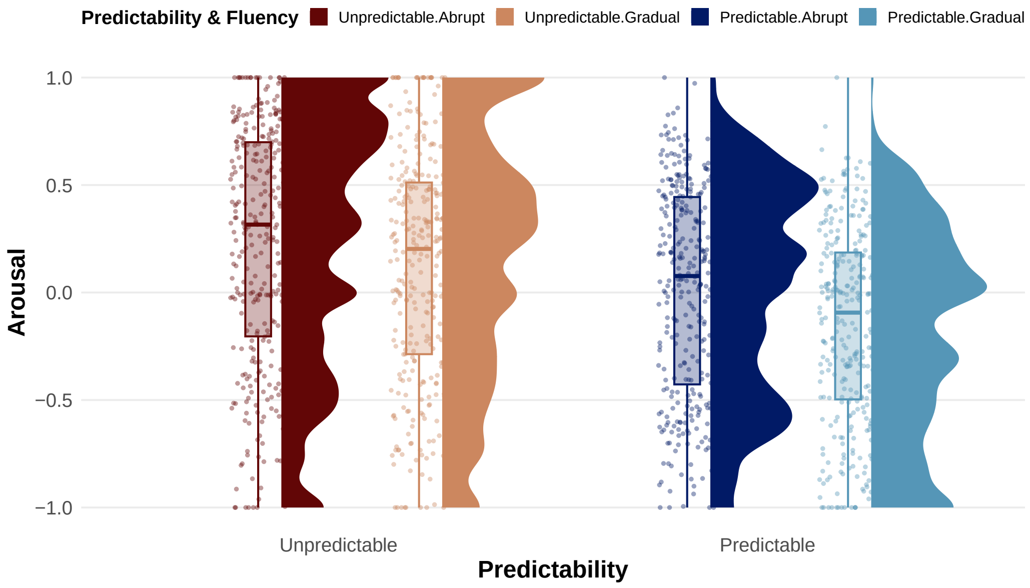


Figure S4: Raincloud plots of subjective arousal ratings. The x-axis displays the combinations of each level of the conditions. The y-axis represents the arousal rating on a continuous scale from -1 (most negative) to 1 (most positive).

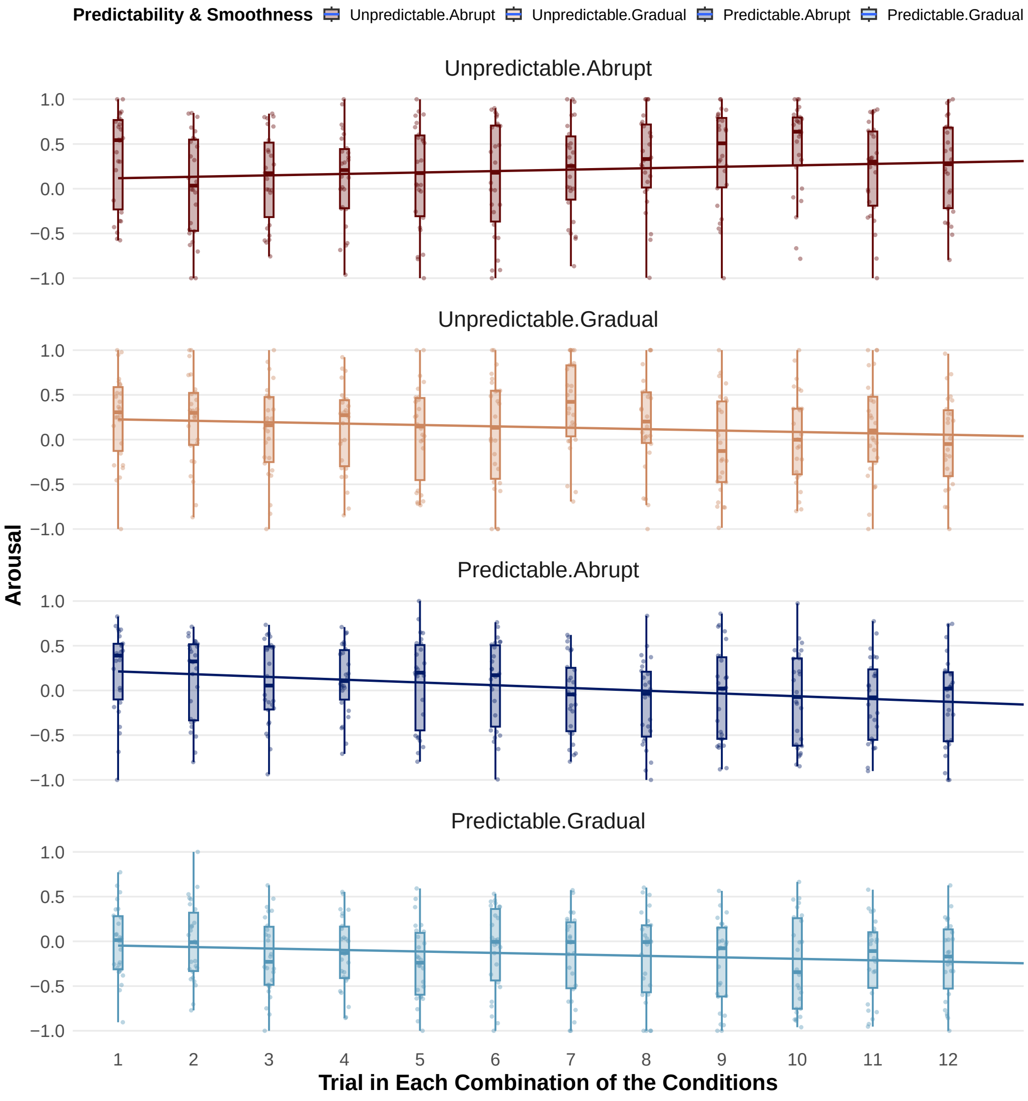


Figure S5: Scattered plots with box plots and regression lines of subjective ratings of arousal in each combination of the conditions. The x-axis displays the trials in a block associated with each combination of the conditions. The y-axis represents the arousal rating.

The mean and standard error of arousal rating for each condition were as follows: `Unpredictable-Abrupt`: $0.221 \pm 0.031$, `Unpredictable-Gradual`: $0.124 \pm 0.030$, `Predictable-Abrupt`: $0.012 \pm 0.028$, `Predictable-Gradual`:

-0.154 ± 0.026. For the summary for each of the four conditions throughout the trials, see Supplement D.

#### *S.2.3. Trust*

Figure S6 and S7 depicts the distribution of trust scored by the participants for each of the four conditions and the distribution throughout the trials in each condition, respectively.

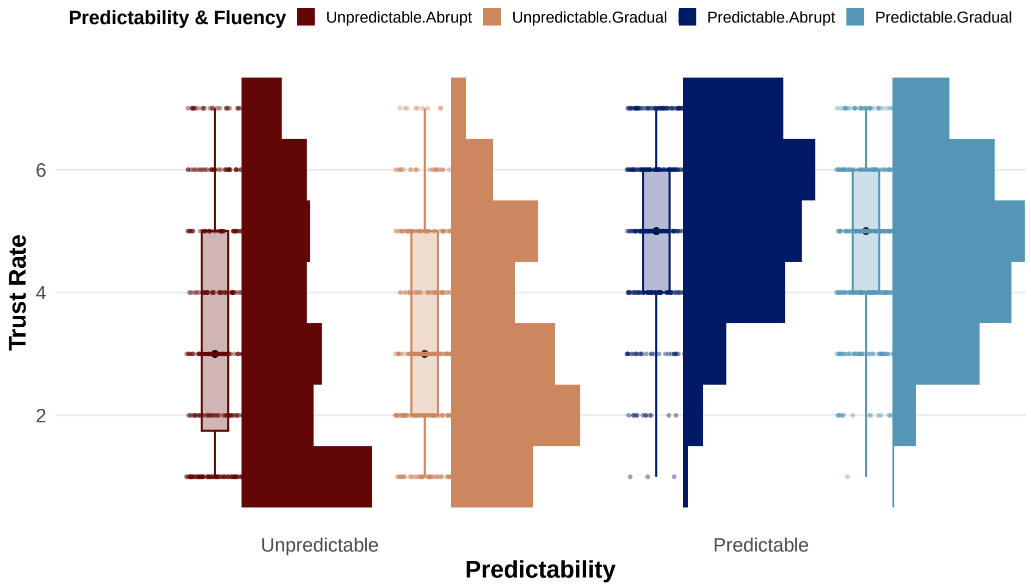


Figure S6: Raincloud plots of subjective trust ratings. The x-axis displays the combinations of each level of the conditions. The y-axis represents the trust rating on a discrete scale from 1 (most negative) to 7 (most positive).

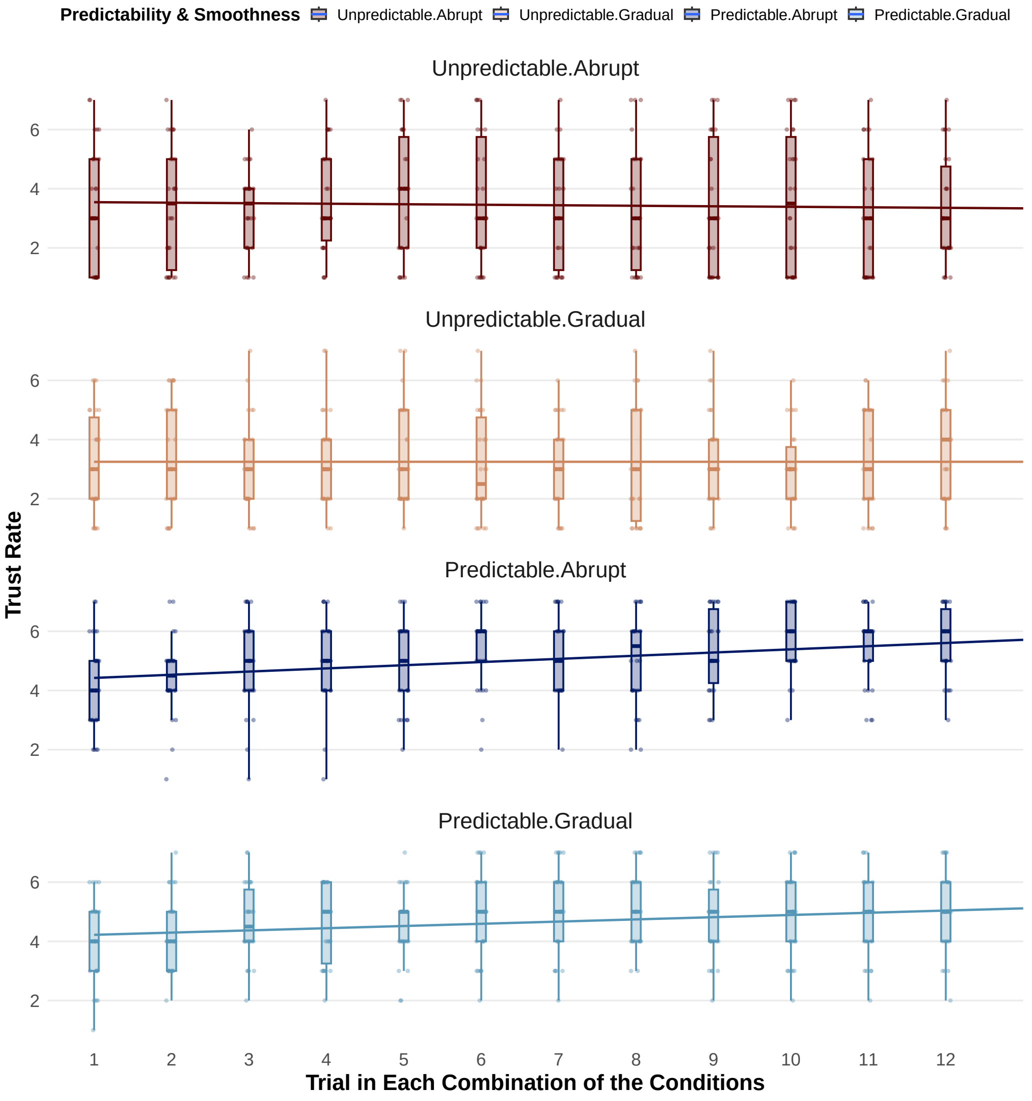


Figure S7: Scattered plots with box plots and regression lines of subjective ratings of trust rate in each combination of the conditions. The x-axis displays the trials in a block associated with each combination of the conditions. The y-axis represents the trust rating.

The mean and standard error of trust rating for each condition were as follows: `Unpredictable-Abrupt`: $3.433 \pm 0.113$, `Unpredictable-Gradual`: $3.250 \pm 0.095$, `Predictable-Abrupt`: $5.122 \pm 0.081$, `Predictable-Gradual`:

4.705 ± 0.078. or the summary for each of the four conditions throughout the trials, see Supplement D.

#### *S.2.4. Skin Conductance Response*

Figure S8 shows the time series data of SCR across all participants, with each trial time normalized to the range of 0% to 100%. The data is grouped by the combination of each level of predictability and smoothness conditions.

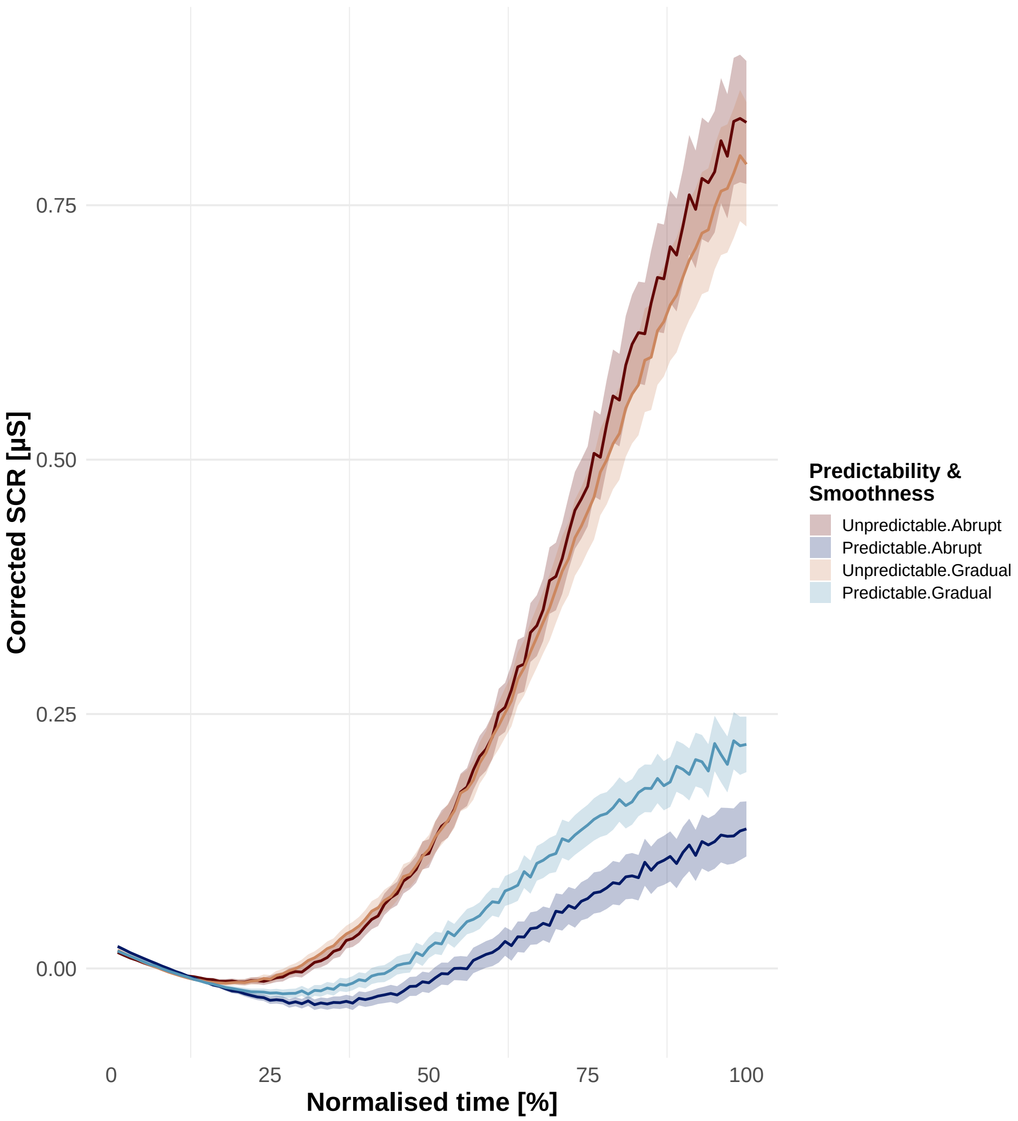


Figure S8: Time series data of SCR. The x-axis displays the trial time normalized to 0%–100%. The y-axis represents the SCR amplitude.

Figure S9 and S10 depicts the raincloud plots of the peak of SCR for each of the four conditions and the distribution throughout the trials in each condition, respectively.

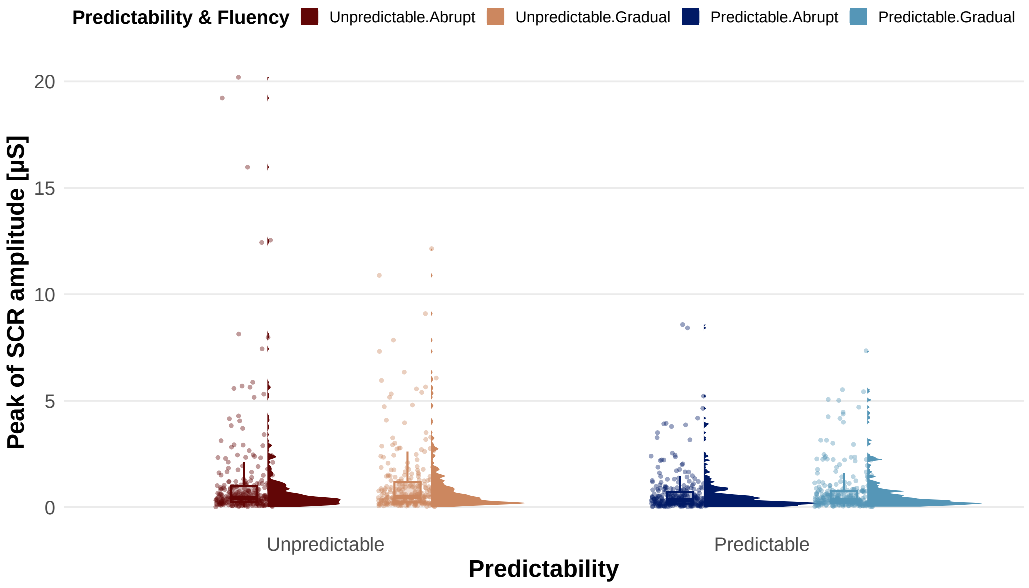


Figure S9: Raincloud plots of SCR. The x-axis displays the combinations of each level of the conditions. The y-axis represents the peak of SCR amplitude.

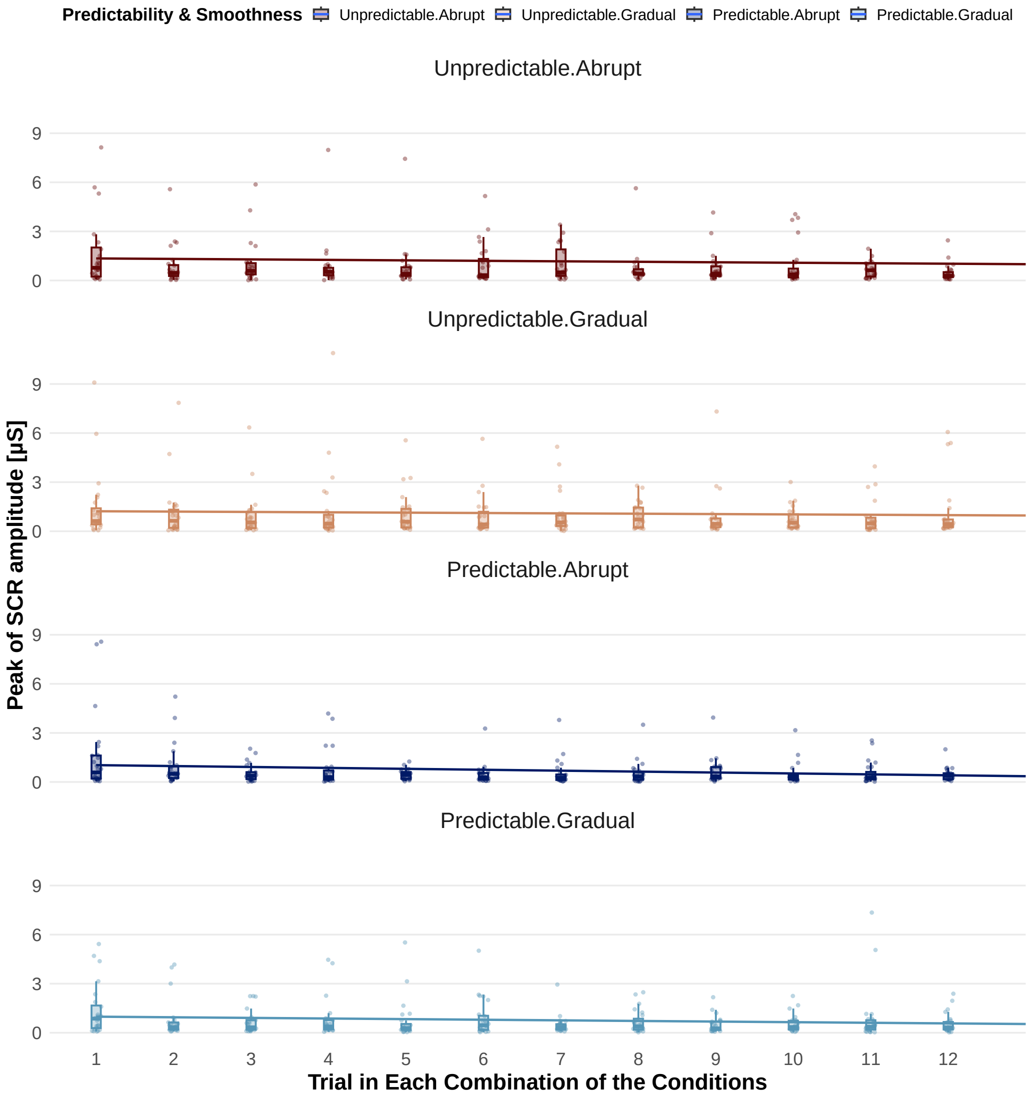


Figure S10: Scattered plots with box plots and regression lines of the peaks of SCR amplitude in each combination of the conditions. The x-axis displays the trials in a block associated with each combination of the conditions. The y-axis represents the peak of SCR amplitude.

The mean and standard error of the peak of SCR amplitude for each condition were as follows: `Unpredictable-Abrupt`: $1.156 \pm 0.141$, `Unpredictable-Gradual`: $1.082 \pm 0.096$, `Predictable-Abrupt`: $0.664 \pm 0.061$, `Predictable-Gradual`:

0.740 ± 0.062.

Figure S11 shows the raincloud plots of the log-transformed peak of SCR amplitude across conditions.

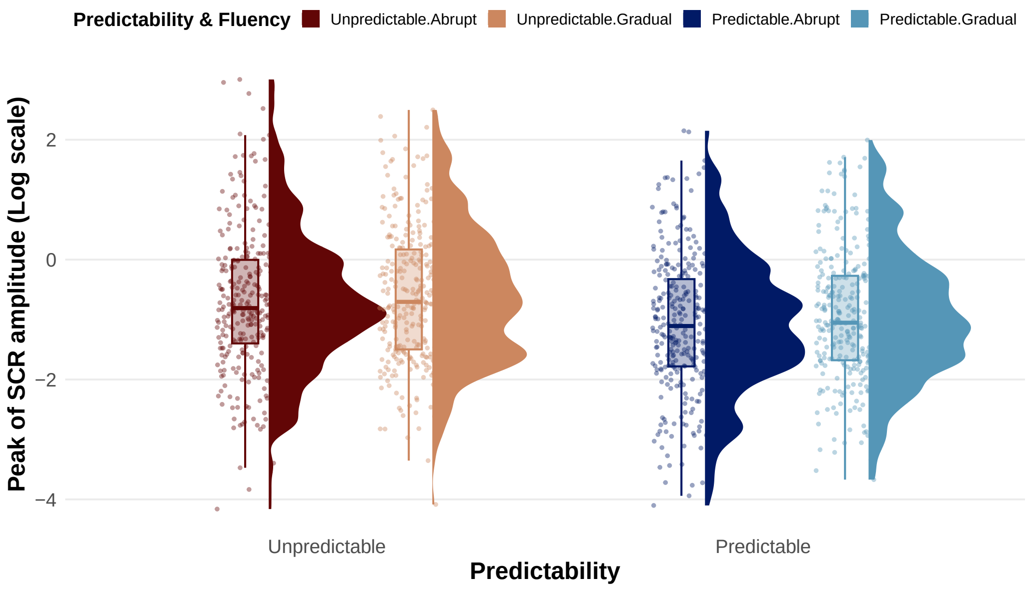


Figure S11: Raincloud plots of log-transformed peak of SCR. The y-axis represents the log-scaled peak of SCR amplitude. The x-axis displays the combinations of each level of the conditions.

For the correlation between SCR and arousal, we fitted a LMM to predict log-scaled SCR amplitude using mean-centered subjective arousal rating as a fixed effect, and by-participant random intercepts and slopes. The fixed effect of the mean-centered arousal was statistically significant (Estimate = 0.334, SE = 0.075, $t(19.723) = 4.454$, $p$ <0.001). The marginal $R^2$ indicating variance explained by fixed effects only was and 0.022, and the conditional $R^2$ indicating variance explained by both fixed and random effects was 0.439). These values show that while the overall fixed effect accounted for approximately 2% of the variance in the peak of SCR amplitude, while the full model including individual differences explained approximately 44%.

Figure S12 and S13 depicts the correlation between the log-scaled peak of SCR amplitude and the mean-centered arousal rating and the distribution of participant-specific slopes for the effect of the arousal rating on the log-scaled peak of SCR amplitude for each participant, respectively.

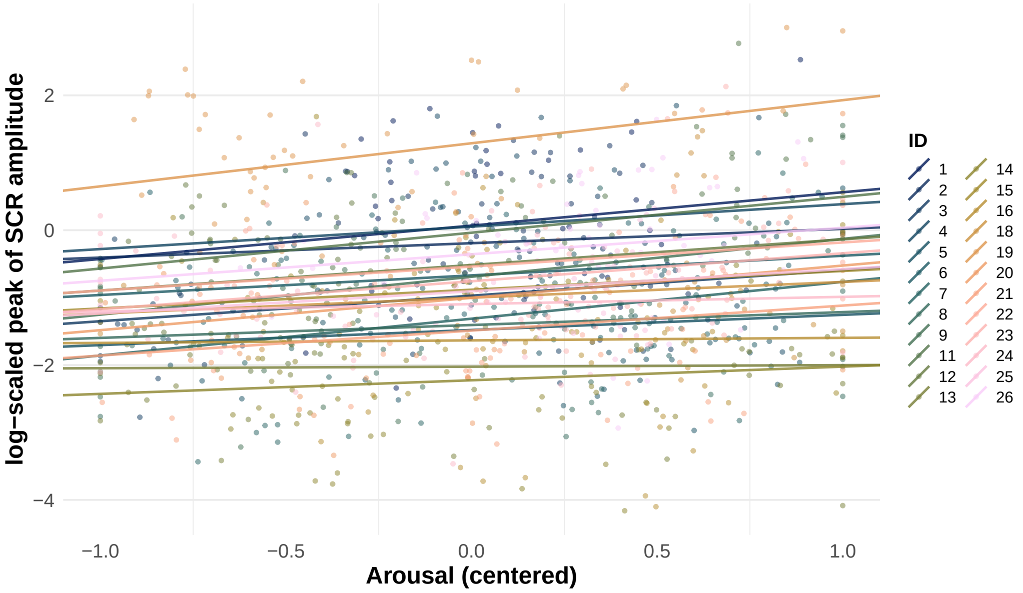


Figure S12: Scatterplot showing the correlation between the log-scaled peak of SCR amplitude and mean-centered arousal rating. Each dot represents the data points for each trial. The X-axis indicates centered arousal ratings. The Y-axis represents the peak SCR amplitude on a logarithmic scale. IDs 10 and 17 were excluded based on the criteria.

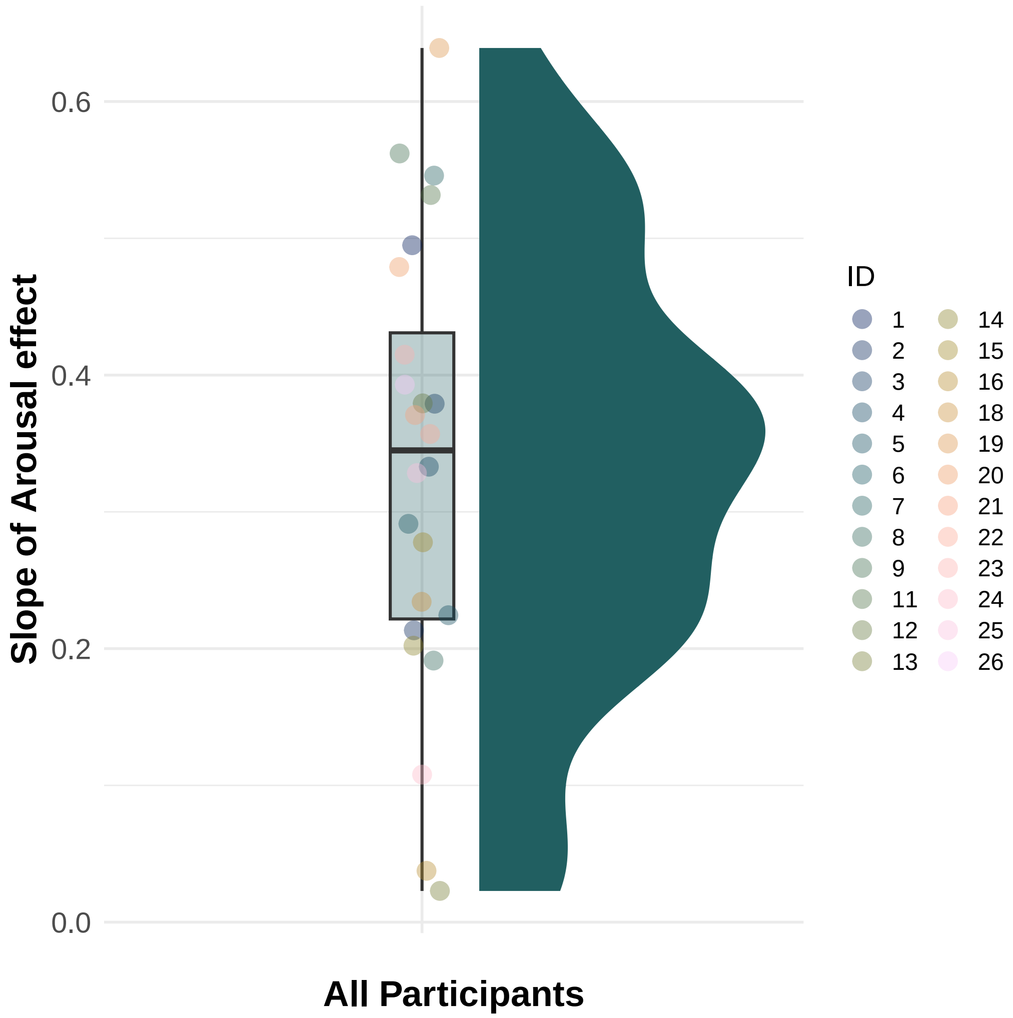


Figure S13: Raincloud plot showing the distribution of participant-specific slopes for the effect of arousal on log-transformed SCR amplitude. The x-axis is used only for plotting all data in one group. The y-axis shows represents individual slope estimated using the GLMM model. The y-axis Each dot represents an individual participant ' s estimated slope from the mixed-effects model.

### *S.3. Diagnostic Checks for the Mixed-Effects Models*

#### *S.3.1. Valence*

For the final model structure for valence (`Valence ~ Predictability * Smoothness * Trial_in_Block_centered + (1 | ID)`), diagnostic checks shown in Figure S14 indicated no substantial violations of model assumptions.

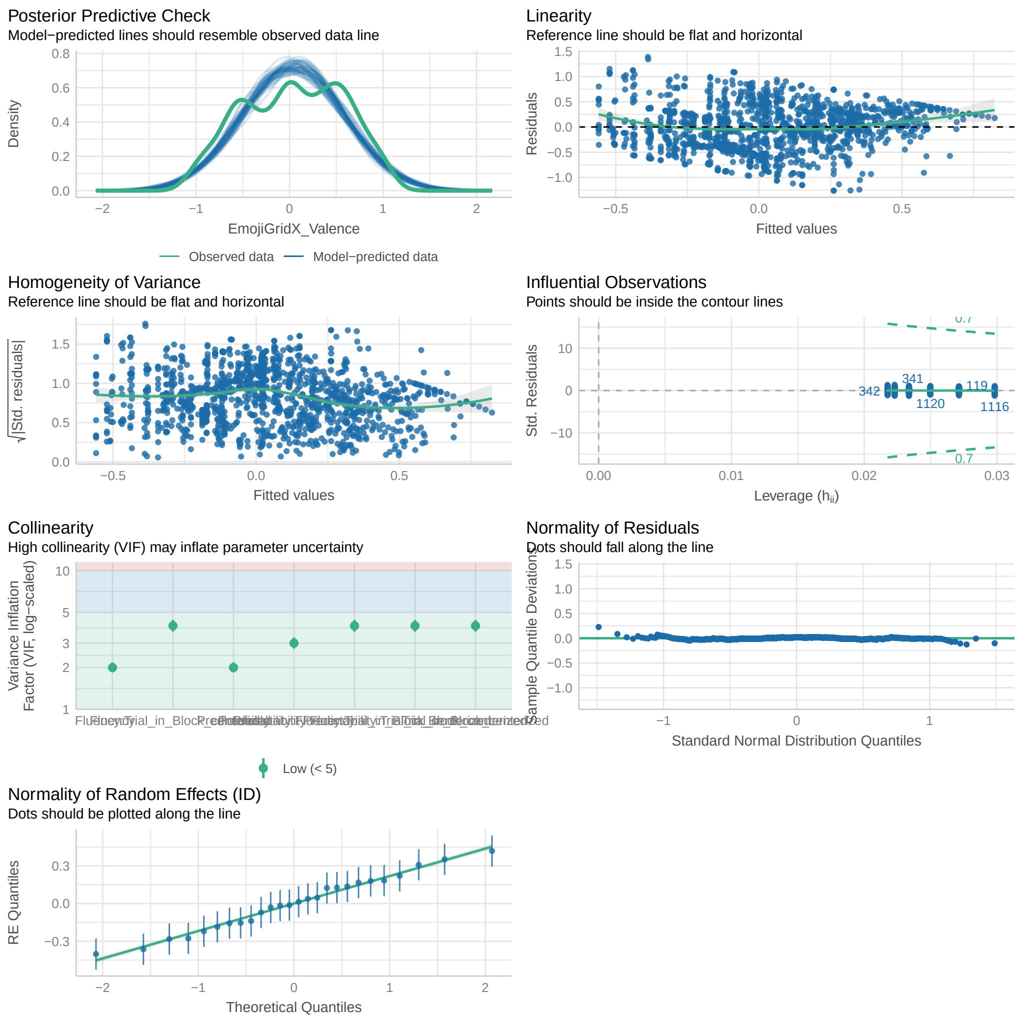


Figure S14: Model diagnostics for the best model of valence

### *S.3.2. Arousal*

For the final model structure for arousal (`Arousal ~ Predictability * Smoothness * Trial_in_Block_centered + (1 + Trial_in_Block_centered || ID)`), the optimizer indicate that the model fit was stable, although the model fit produced a convergence warning. Diagnostic checks shown in Figure S15 indicated no other critical violations of the assumptions.

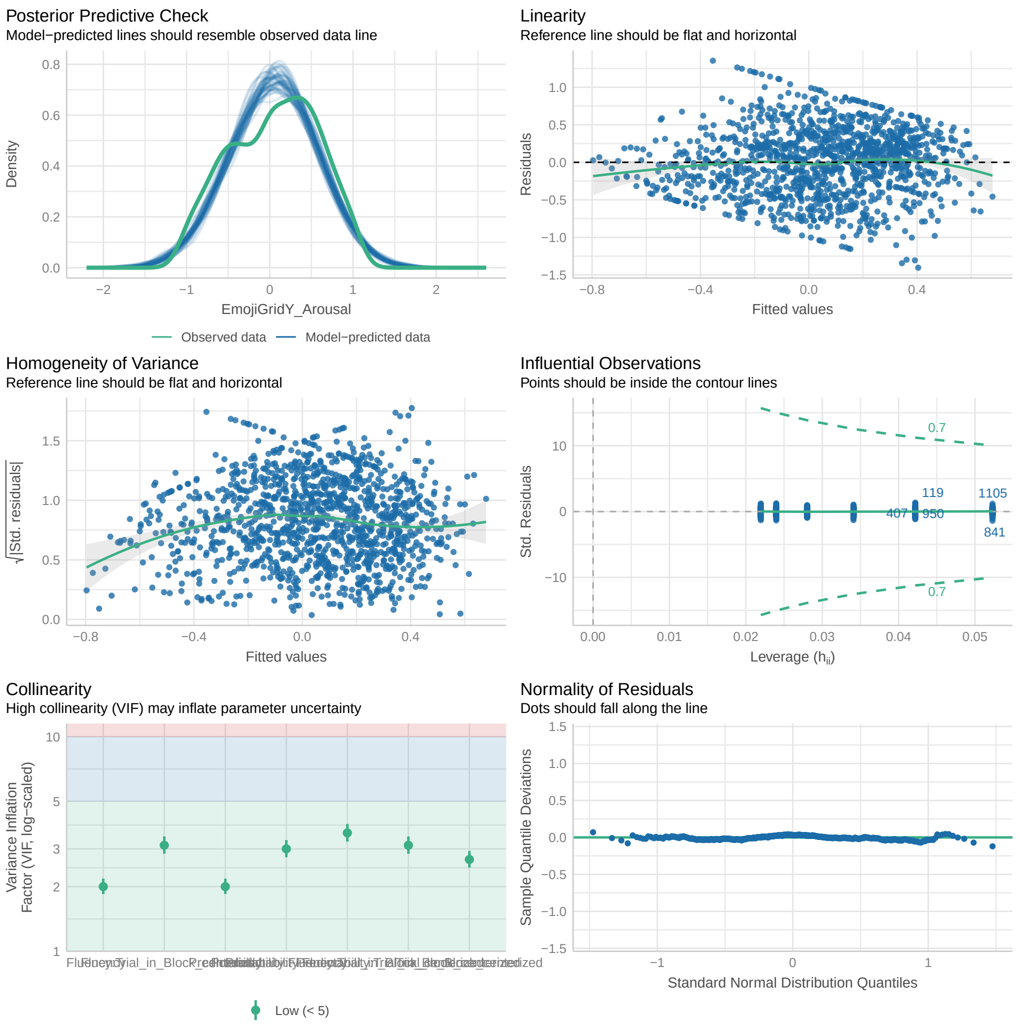


Figure S15: Model diagnostics for the best model of arousal

#### *S.3.3. Skin Conductance Response*

For the LLM structure for skin conductance response, diagnostic checks shown in Figure S16 indicated substantial violations of the assumptions, particularly regarding the homogeneity of variance and the normality of residuals.

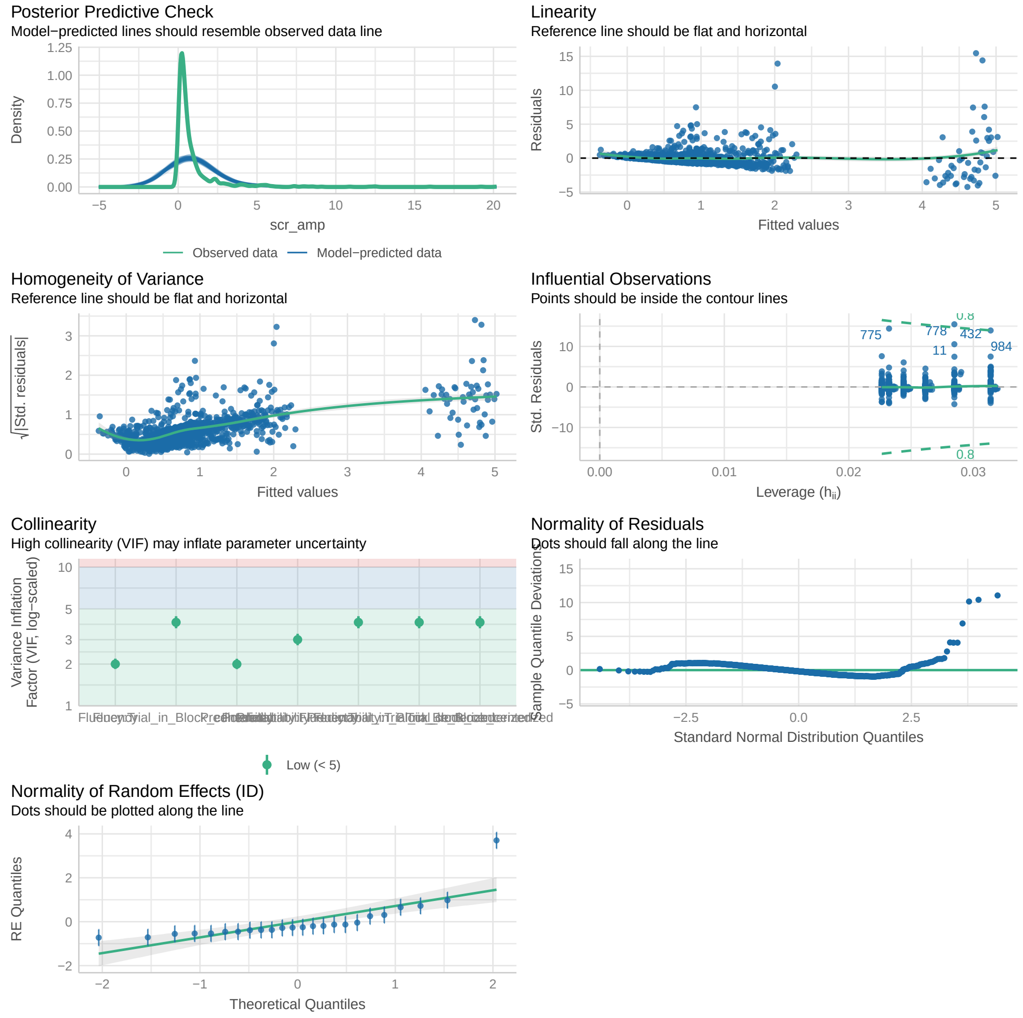


Figure S16: Model diagnostics for the best linear mixed model of the peak of SCR

For the GLMM structure for skin conductance response that was built to address the violations of the assumptions, (`SCRamplitude ~ Predictability * Smoothness * Trial_in_Block_centered + (1 + Trial_in_Block_centered | ID)`), diagnostic checks shown in Figure S17 indicated no substantial violations of the assumptions.

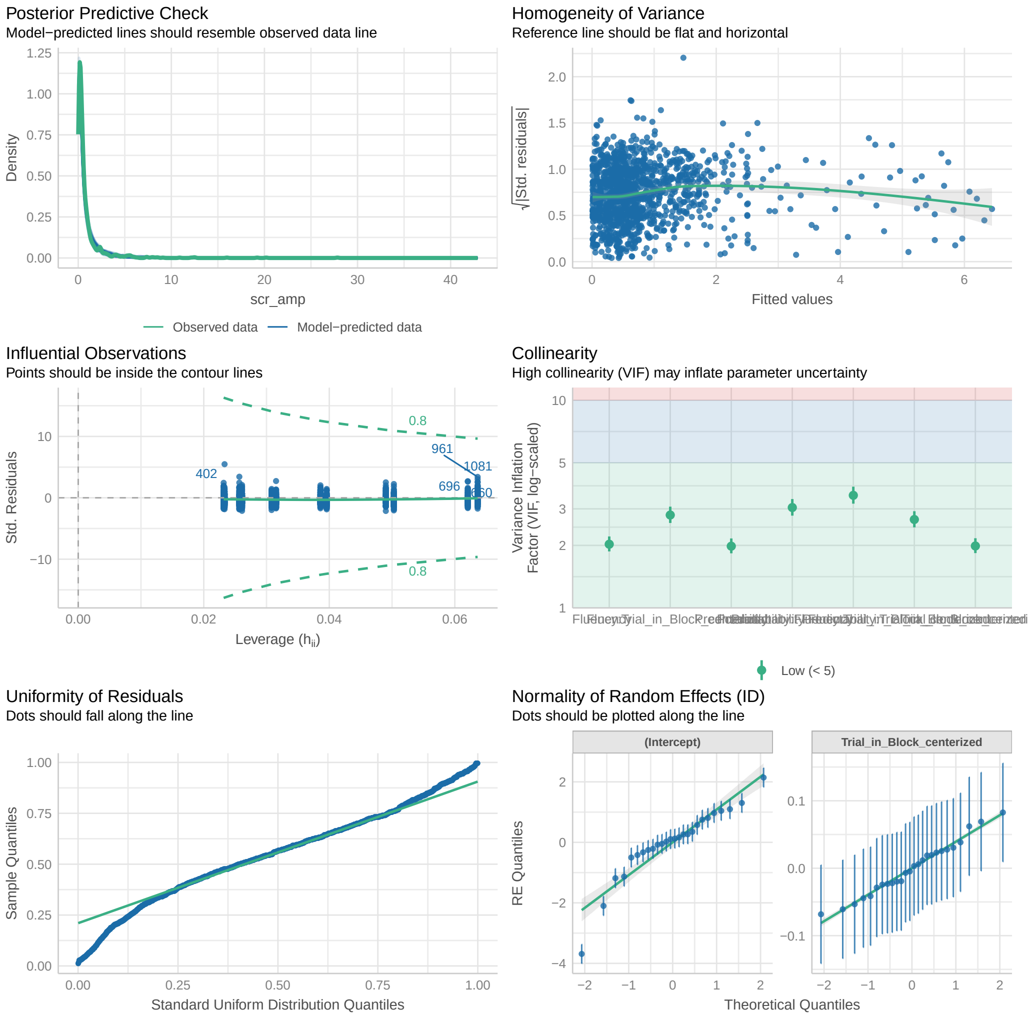


Figure S17: Model diagnostics for the best generarized linear mixed model of the peak of SCR

### *S.4. Summary Table of the Results*

Table 9: Summary of subjective ratings of valence across conditions and trials.

| Predictability | Smoothness | Trial in Block | Mean Valence | SE Valence |
|---|---|---|---|---|
| Unpredictable | Abrupt | 1 | -0.148 | 0.103 |
| Unpredictable | Abrupt | 2 | -0.109 | 0.112 |

| | | | | |
|---|---|---|---|---|
| Unpredictable | Abrupt | 3 | -0.123 | 0.095 |
| Unpredictable | Abrupt | 4 | -0.112 | 0.095 |
| Unpredictable | Abrupt | 5 | 0.048 | 0.108 |
| Unpredictable | Abrupt | 6 | -0.133 | 0.108 |
| Unpredictable | Abrupt | 7 | -0.163 | 0.099 |
| Unpredictable | Abrupt | 8 | -0.123 | 0.104 |
| Unpredictable | Abrupt | 9 | -0.058 | 0.117 |
| Unpredictable | Abrupt | 10 | -0.085 | 0.125 |
| Unpredictable | Abrupt | 11 | -0.123 | 0.108 |
| Unpredictable | Abrupt | 12 | -0.113 | 0.099 |
| Unpredictable | Gradual | 1 | -0.219 | 0.097 |
| Unpredictable | Gradual | 2 | -0.084 | 0.098 |
| Unpredictable | Gradual | 3 | -0.245 | 0.101 |
| Unpredictable | Gradual | 4 | -0.065 | 0.106 |
| Unpredictable | Gradual | 5 | -0.019 | 0.125 |
| Unpredictable | Gradual | 6 | -0.163 | 0.107 |
| Unpredictable | Gradual | 7 | -0.240 | 0.091 |
| Unpredictable | Gradual | 8 | -0.318 | 0.112 |
| Unpredictable | Gradual | 9 | -0.080 | 0.105 |
| Unpredictable | Gradual | 10 | -0.242 | 0.088 |
| Unpredictable | Gradual | 11 | -0.083 | 0.114 |
| Unpredictable | Gradual | 12 | -0.128 | 0.106 |
| Predictable | Abrupt | 1 | 0.087 | 0.088 |
| Predictable | Abrupt | 2 | 0.182 | 0.076 |
| Predictable | Abrupt | 3 | 0.235 | 0.090 |
| Predictable | Abrupt | 4 | 0.291 | 0.092 |

| | | | | |
|---|---|---|---|---|
| Predictable | Abrupt | 5 | 0.189 | 0.096 |
| Predictable | Abrupt | 6 | 0.382 | 0.093 |
| Predictable | Abrupt | 7 | 0.285 | 0.104 |
| Predictable | Abrupt | 8 | 0.215 | 0.109 |
| Predictable | Abrupt | 9 | 0.332 | 0.095 |
| Predictable | Abrupt | 10 | 0.434 | 0.092 |
| Predictable | Abrupt | 11 | 0.362 | 0.088 |
| Predictable | Abrupt | 12 | 0.381 | 0.098 |
| Predictable | Gradual | 1 | 0.021 | 0.099 |
| Predictable | Gradual | 2 | 0.106 | 0.104 |
| Predictable | Gradual | 3 | 0.188 | 0.094 |
| Predictable | Gradual | 4 | 0.142 | 0.101 |
| Predictable | Gradual | 5 | 0.177 | 0.085 |
| Predictable | Gradual | 6 | 0.196 | 0.100 |
| Predictable | Gradual | 7 | 0.253 | 0.083 |
| Predictable | Gradual | 8 | 0.262 | 0.081 |
| Predictable | Gradual | 9 | 0.097 | 0.082 |
| Predictable | Gradual | 10 | 0.065 | 0.082 |
| Predictable | Gradual | 11 | 0.194 | 0.086 |
| Predictable | Gradual | 12 | 0.149 | 0.092 |

Table 10: Summary of subjective ratings of arousal across conditions and trials.

| Predictability | Smoothness | Trial in Block | Mean Arousal | SE Arousal |
|---|---|---|---|---|
| 0 | 0 | 1 | 0.332 | 0.106 |
| 0 | 0 | 2 | 0.032 | 0.114 |

| | | | | |
|---|---|---|---|---|
| 0 | 0 | 3 | 0.131 | 0.099 |
| 0 | 0 | 4 | 0.128 | 0.101 |
| 0 | 0 | 5 | 0.140 | 0.116 |
| 0 | 0 | 6 | 0.118 | 0.125 |
| 0 | 0 | 7 | 0.205 | 0.105 |
| 0 | 0 | 8 | 0.314 | 0.103 |
| 0 | 0 | 9 | 0.349 | 0.108 |
| 0 | 0 | 10 | 0.466 | 0.099 |
| 0 | 0 | 11 | 0.198 | 0.104 |
| 0 | 0 | 12 | 0.243 | 0.103 |
| 0 | 1 | 1 | 0.242 | 0.097 |
| 0 | 1 | 2 | 0.217 | 0.098 |
| 0 | 1 | 3 | 0.096 | 0.102 |
| 0 | 1 | 4 | 0.117 | 0.095 |
| 0 | 1 | 5 | 0.086 | 0.106 |
| 0 | 1 | 6 | 0.086 | 0.123 |
| 0 | 1 | 7 | 0.386 | 0.101 |
| 0 | 1 | 8 | 0.182 | 0.107 |
| 0 | 1 | 9 | -0.051 | 0.110 |
| 0 | 1 | 10 | 0.028 | 0.101 |
| 0 | 1 | 11 | 0.128 | 0.110 |
| 0 | 1 | 12 | -0.025 | 0.100 |
| 1 | 0 | 1 | 0.215 | 0.093 |
| 1 | 0 | 2 | 0.134 | 0.092 |
| 1 | 0 | 3 | 0.085 | 0.093 |
| 1 | 0 | 4 | 0.119 | 0.077 |

| | | | | |
|---|---|---|---|---|
| 1 | 0 | 5 | 0.102 | 0.101 |
| 1 | 0 | 6 | 0.071 | 0.095 |
| 1 | 0 | 7 | -0.067 | 0.087 |
| 1 | 0 | 8 | -0.130 | 0.093 |
| 1 | 0 | 9 | -0.062 | 0.105 |
| 1 | 0 | 10 | -0.091 | 0.104 |
| 1 | 0 | 11 | -0.111 | 0.093 |
| 1 | 0 | 12 | -0.115 | 0.100 |
| 1 | 1 | 1 | -0.013 | 0.079 |
| 1 | 1 | 2 | -0.013 | 0.087 |
| 1 | 1 | 3 | -0.167 | 0.085 |
| 1 | 1 | 4 | -0.131 | 0.080 |
| 1 | 1 | 5 | -0.234 | 0.083 |
| 1 | 1 | 6 | -0.084 | 0.093 |
| 1 | 1 | 7 | -0.159 | 0.092 |
| 1 | 1 | 8 | -0.159 | 0.099 |
| 1 | 1 | 9 | -0.250 | 0.095 |
| 1 | 1 | 10 | -0.249 | 0.104 |
| 1 | 1 | 11 | -0.191 | 0.087 |
| 1 | 1 | 12 | -0.202 | 0.086 |

Table 11: Summary of subjective ratings of trust across conditions and trials.

| Predictability | Smoothness | Trial in Block | Mean Rate of Trust | SE Rate of Trust |
|---|---|---|---|---|
| 0 | 0 | 1 | 3.346 | 0.415 |
| 0 | 0 | 2 | 3.462 | 0.397 |

| | | | | |
|---|---|---|---|---|
| 0 | 0 | 3 | 3.269 | 0.280 |
| 0 | 0 | 4 | 3.731 | 0.353 |
| 0 | 0 | 5 | 3.808 | 0.404 |
| 0 | 0 | 6 | 3.577 | 0.430 |
| 0 | 0 | 7 | 3.308 | 0.371 |
| 0 | 0 | 8 | 3.308 | 0.411 |
| 0 | 0 | 9 | 3.385 | 0.437 |
| 0 | 0 | 10 | 3.615 | 0.454 |
| 0 | 0 | 11 | 3.077 | 0.407 |
| 0 | 0 | 12 | 3.308 | 0.358 |
| 0 | 1 | 1 | 3.154 | 0.336 |
| 0 | 1 | 2 | 3.538 | 0.369 |
| 0 | 1 | 3 | 3.077 | 0.313 |
| 0 | 1 | 4 | 3.385 | 0.314 |
| 0 | 1 | 5 | 3.385 | 0.338 |
| 0 | 1 | 6 | 3.115 | 0.352 |
| 0 | 1 | 7 | 3.077 | 0.298 |
| 0 | 1 | 8 | 3.154 | 0.383 |
| 0 | 1 | 9 | 3.385 | 0.314 |
| 0 | 1 | 10 | 2.962 | 0.274 |
| 0 | 1 | 11 | 3.192 | 0.333 |
| 0 | 1 | 12 | 3.577 | 0.338 |
| 1 | 0 | 1 | 4.269 | 0.302 |
| 1 | 0 | 2 | 4.462 | 0.262 |
| 1 | 0 | 3 | 5.000 | 0.319 |
| 1 | 0 | 4 | 5.000 | 0.283 |

| | | | | |
|---|---|---|---|---|
| 1 | 0 | 5 | 4.885 | 0.290 |
| 1 | 0 | 6 | 5.423 | 0.255 |
| 1 | 0 | 7 | 5.269 | 0.263 |
| 1 | 0 | 8 | 5.038 | 0.311 |
| 1 | 0 | 9 | 5.346 | 0.266 |
| 1 | 0 | 10 | 5.692 | 0.220 |
| 1 | 0 | 11 | 5.462 | 0.256 |
| 1 | 0 | 12 | 5.615 | 0.229 |
| 1 | 1 | 1 | 4.000 | 0.272 |
| 1 | 1 | 2 | 4.192 | 0.248 |
| 1 | 1 | 3 | 4.615 | 0.255 |
| 1 | 1 | 4 | 4.577 | 0.249 |
| 1 | 1 | 5 | 4.577 | 0.243 |
| 1 | 1 | 6 | 4.846 | 0.287 |
| 1 | 1 | 7 | 5.077 | 0.288 |
| 1 | 1 | 8 | 5.154 | 0.252 |
| 1 | 1 | 9 | 4.731 | 0.275 |
| 1 | 1 | 10 | 4.846 | 0.292 |
| 1 | 1 | 11 | 4.808 | 0.266 |
| 1 | 1 | 12 | 5.038 | 0.274 |

Table 12: Summary of subjective ratings of arousal across conditions and trials.

| Predictability | Smoothness | Trial in Block | Mean Peak of SCR | SE Peak of SCR |
|---|---|---|---|---|
| 0 | 0 | 1 | 1.986 | 0.685 |
| 0 | 0 | 2 | 0.820 | 0.232 |

| | | | | |
|---|---|---|---|---|
| 0 | 0 | 3 | 0.987 | 0.265 |
| 0 | 0 | 4 | 0.829 | 0.299 |
| 0 | 0 | 5 | 0.790 | 0.279 |
| 0 | 0 | 6 | 0.950 | 0.241 |
| 0 | 0 | 7 | 1.338 | 0.485 |
| 0 | 0 | 8 | 1.386 | 0.743 |
| 0 | 0 | 9 | 0.692 | 0.182 |
| 0 | 0 | 10 | 0.855 | 0.245 |
| 0 | 0 | 11 | 1.794 | 0.873 |
| 0 | 0 | 12 | 0.439 | 0.103 |
| 0 | 1 | 1 | 1.322 | 0.396 |
| 0 | 1 | 2 | 1.073 | 0.333 |
| 0 | 1 | 3 | 0.892 | 0.262 |
| 0 | 1 | 4 | 1.235 | 0.447 |
| 0 | 1 | 5 | 1.018 | 0.251 |
| 0 | 1 | 6 | 1.902 | 1.064 |
| 0 | 1 | 7 | 0.953 | 0.250 |
| 0 | 1 | 8 | 0.883 | 0.155 |
| 0 | 1 | 9 | 0.848 | 0.291 |
| 0 | 1 | 10 | 0.692 | 0.139 |
| 0 | 1 | 11 | 1.180 | 0.480 |
| 0 | 1 | 12 | 1.054 | 0.338 |
| 1 | 0 | 1 | 1.432 | 0.456 |
| 1 | 0 | 2 | 0.872 | 0.243 |
| 1 | 0 | 3 | 0.494 | 0.107 |
| 1 | 0 | 4 | 0.736 | 0.221 |

| | | | | |
|---|---|---|---|---|
| 1 | 0 | 5 | 0.409 | 0.062 |
| 1 | 0 | 6 | 0.440 | 0.124 |
| 1 | 0 | 7 | 0.497 | 0.156 |
| 1 | 0 | 8 | 0.483 | 0.140 |
| 1 | 0 | 9 | 0.612 | 0.156 |
| 1 | 0 | 10 | 0.443 | 0.131 |
| 1 | 0 | 11 | 0.522 | 0.131 |
| 1 | 0 | 12 | 0.424 | 0.080 |
| 1 | 1 | 1 | 1.249 | 0.300 |
| 1 | 1 | 2 | 0.735 | 0.223 |
| 1 | 1 | 3 | 0.600 | 0.135 |
| 1 | 1 | 4 | 0.757 | 0.227 |
| 1 | 1 | 5 | 0.673 | 0.234 |
| 1 | 1 | 6 | 0.832 | 0.220 |
| 1 | 1 | 7 | 0.449 | 0.111 |
| 1 | 1 | 8 | 0.618 | 0.137 |
| 1 | 1 | 9 | 0.457 | 0.098 |
| 1 | 1 | 10 | 0.506 | 0.108 |
| 1 | 1 | 11 | 0.841 | 0.323 |
| 1 | 1 | 12 | 0.508 | 0.119 |
